# Competitive fungal commensalism mitigates candidiasis pathology

**DOI:** 10.1101/2024.01.12.575358

**Authors:** Jarmila Sekeresova Kralova, Catalina Donic, Bareket Dassa, Ilana Livyatan, Paul Mathias Jansen, Shifra Ben-Dor, Lena Fidel, Sébastien Trzebanski, Lian Narunsky-Haziza, Omer Asraf, Ori Brenner, Hagit Dafni, Ghil Jona, Sigalit Boura-Halfon, Noa Stettner, Eran Segal, Sascha Brunke, Yitzhak Pilpel, Ravid Straussman, David Zeevi, Petra Bacher, Bernhard Hube, Neta Shlezinger, Steffen Jung

## Abstract

The mycobiota are a critical part of the gut microbiome, but host-fungal interactions and specific functional contributions of commensal fungi to host fitness remain incompletely understood. Here we report the identification of a new fungal commensal, *Kazachstania heterogenica* var. *weizmannii,* isolated from murine intestines. *K. weizmannii* exposure prevented *Candida albicans* colonization and significantly reduced the commensal *C. albicans* burden in colonized animals. Following immunosuppression of *C. albicans* colonized mice, competitive fungal commensalism thereby mitigated fatal candidiasis. Metagenome analysis revealed *K. weizmannii* presence among human commensals. Our results reveal competitive fungal commensalism within the intestinal microbiota, independent of bacteria and immune responses, that could bear potential therapeutic value for the management of *C. albicans*-mediated diseases.

## Introduction

The genetic information that defines human beings includes the genomes of the human cells but also that of the commensal microbiome. While the analysis of this metagenome has focused mainly on the abundant bacterial commensals ^1,2^, mucosal surfaces are also inhabited by fungi. The role of the mycobiota and their contributions to host fitness are, however, less well understood ^3,4^. This is in part due to the underrepresentation of fungi in the microbiome of animal models that are kept under strict hygienic conditions ^5–7^.

As one of the most common human fungal pathogens, *Candida albicans* causes hundreds of millions of symptomatic infections each year ^8,9^. Pathologies are frequently associated with immuno-deficiencies and range from superficial irritations of the skin and mucosae to life-threatening invasive infections of internal organs. In addition, chronic mucocutaneous candidiasis (CMC) has been linked to inborn errors of IL-17 immunity ^10^. Fungal dissemination that leads to systemic infection, is believed to originate from the gut, where *C. albicans* normally resides as a harmless commensal ^11^. Candidiasis has been linked to filamentation of the fungus ^12,13^, which is strictly associated with the expression of the cytolytic peptide toxin candidalysin that promotes barrier damage ^14,15^. Human individuals are colonized with *C. albicans* in childhood, and clonal fungal populations persist over their lifetime, mostly without symptoms. Emerging evidence suggests that fungal colonization in fact rather benefits, than harms the host and that the gut mycobiota improve mammalian immunity ^16–19^. Specifically, commensal fungi, and in particular *C. albicans*, were shown to affect the composition of the myeloid innate immune compartment ^5,20,21^ and elicit cellular and humoral immunity ^22–24^. Insights into the diverse interactions of fungi with the mammalian hosts and other fungi, but also communication with bacterial commensals, could aid our understanding of host physiology. Moreover, a better understanding could allow harnessing the impact of commensal fungi on human immunity for therapeutic purposes.

The host-fungi interface remains, however, incompletely understood, not the least due to the lack of suitable experimental animal models and our limited insight into fungal commensalism.

The dominant human fungal commensal, *C. albicans*, has also been reported as part of the mycobiota of mice roaming in the wild ^5,25^. However, animals kept under specific pathogen-free (SPF) conditions mostly lack *C. albicans* and generally harbor poorly developed mycobiota that also differ considerably between vendors ^26^. Indeed, laboratory mice largely resist *C. albicans* colonization unless subjected to antibiotics (Abx) ^27^, which neutralize inhibiting bacteria, including Lactobacillae ^28,29^.

Here we report the serendipitous identification of a novel fungal commensal of the *Kazachstania* genus that efficiently colonizes laboratory animals kept in SPF facilities without prior Abx conditioning. The strain, which we termed *Kazachstania heterogenica* var. *weizmannii* (*K. weizmannii)*, prevented *C. albicans* colonization, outcompeted *C. albicans* during competitive seeding and even expelled *C. albicans* from stably colonized animals. Murine hosts mounted comparable humoral, but distinct cellular immune responses against *K. weizmannii* and *C. albicans*, although the latter were not required for the competition phenomenon. Unlike *C. albicans*, non-filamenting *K. weizmannii* did not disseminate or cause pathology in immunosuppressed animals. Rather, inter-fungal competition that reduced the intestinal *C. albicans* load of mice, mitigated fatal candidiasis. Finally, human metagenome analysis revealed *K. weizmannii* as a component of the commensal intestinal and vaginal microbiota. Taken together we report the identification of a new fungus that shows robust competitive commensalism with *C. albicans* in mice and thereby mitigating systemic fungal pathology.

## Results

### Identification of a novel commensal fungus in laboratory mice

To probe for a role of myeloid cells in anti-fungal immunity, we generated mice that harbor a corresponding IL-23 deficiency and attempted to colonize them with *C. albicans*. Specifically, we used a protocol involving prior conditioning of animals with antibiotics (Abx) ^27^ (**Fig. 1A**). Ampicillin-exposed wildtype (WT) mice could be readily and persistently seeded with *C. albicans* SC5314 harboring a GFP reporter ^30^ (**Fig. 1B**). However, we consistently failed to efficiently colonize the *Il23a^Δ/Δ^* mice with *C. albicans* (**Fig. 1C**). Following plating of the fecal microbiota of these animals, we noted growth of another yeast-like fungus of distinct morphology (**Fig. 1D, E**). Sequencing of DNA isolated from the plated fungus using the internal transcribed spacer (ITS) as yeast barcode ^31^, tentatively identified the fungus as a member of the *Kazachstania* clade (**Suppl. Fig 1A, B**). This genus is composed of over 50 species found in both anthropic and non-anthropic environments and associated with sourdough production ^32,33^. Whole genome sequencing of the new fungus revealed the characteristic gene duplications of the Saccharomycetaceae family and established it as a novel *Kazachstania* strain. Phylogenetic analysis of 26S rDNA confirmed the assignment of the fungus to the Saccharomycetaceae clade (**Fig. 1F**), and comparison to other *Kazachstania* species (**Fig. 1G**) placed it together with *K. heterogenica*, a fungus found in rodent feces, and in a sister clade with *K sp. Y4206*, a species isolated from human feces. We decided to term the new fungus *Kazachstania heterogenica* var. *weizmannii*, or *K. weizmannii* for short (*Kralova, Fidel et al. submitted*). Comparison of the *K. weizmannii* genome with other full length *Kazachstania* genomes (**Fig. 1H**) showed that *K. heterogenica* is the closest species, followed by *K. pintolopesii, K. telluris* and *K. bovina* in the *Kazachstania telluris* complex (*Ben-Dor, Fidel et al. submitted*). Of note, although we originally discovered the novel *Kazachstania* species in *Il23a^Δ/Δ^* mice that could have impaired anti-fungal immunity, sentinel screening in our animal facilities with the ITS assay revealed that *K. weizmannii* was widespread, irrespective of genotypes of the animals (**Suppl. Fig 1C**).

**Figure 1:**
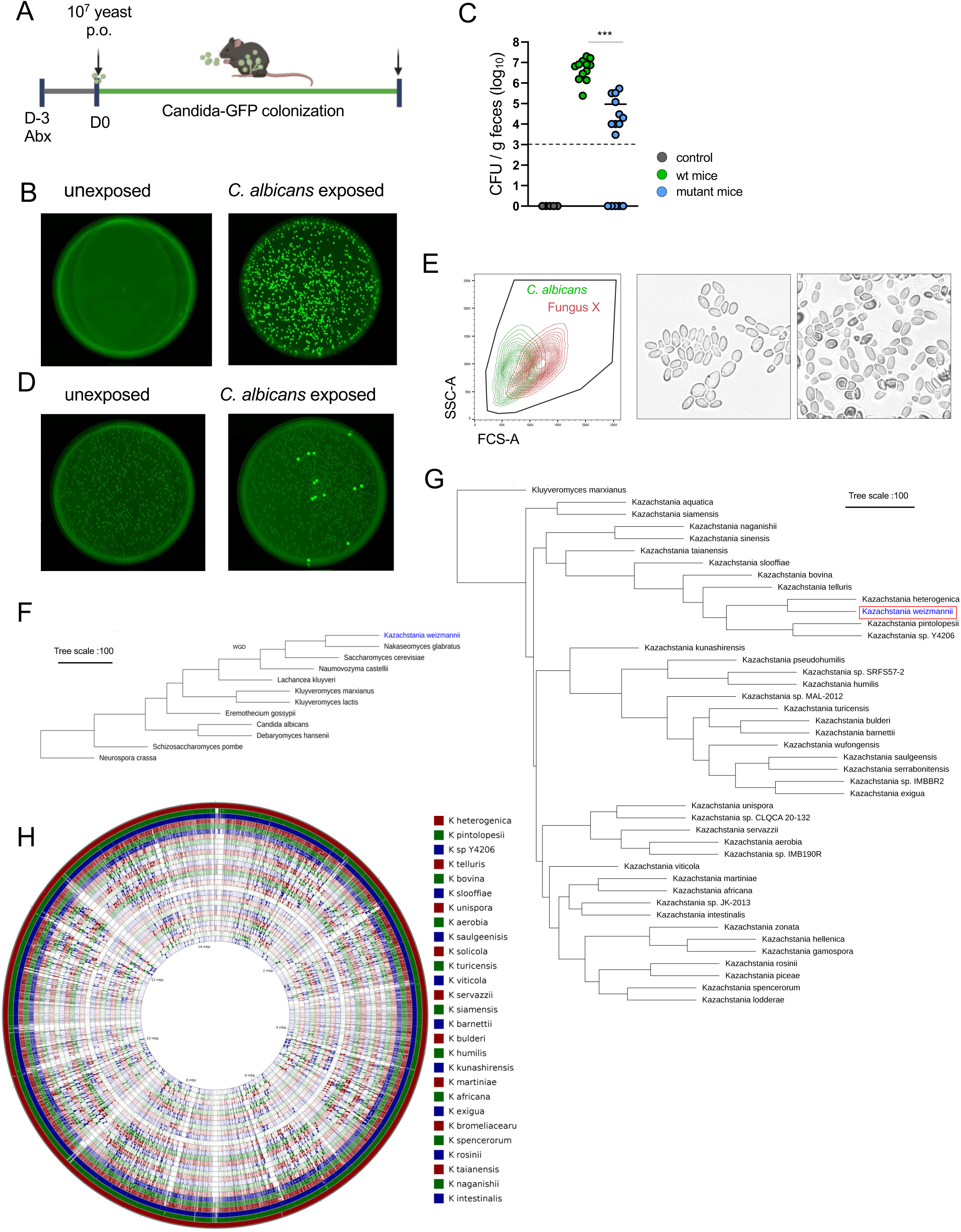
Identification and characterization of *K. weizmannii*. (A) Schematic of *C. albicans* colonization protocol, adopted from (Basu et al., 2000). (B) Recoverable *C. albicans* in the feces of wt mice with Abx supplementation in the drinking water, 2 weeks after oral *C. albicans* inoculation. (C) Recoverable *C. albicans-GFP colonies* in feces of unexposed wt (controls), *C. albicans* inoculated mutant mice and wt animals; stippled line, limit of detection, t test=0.0001 wt vs. mutant Candida colonization. (D) Recoverable *C. albicans-GFP* in the feces 2 weeks after oral inoculation – comparison of unexposed and *Candida*-inoculated mutant animals (E) Flow cytometry and microscopical examination of fungal colonies recovered from mutant animals. (F) Phylogenetic tree based on Maximum Parsimony of the 26S rDNA domains 1 and 2 (D1/D2) region of newly identified *K. weizmannii* in comparison to other yeast model species (G) Phylogenetic tree based on Maximum Parsimony of the 26S rDNA D1/D2 region of *K. weizmannii* in comparison to other *Kazachstania* species. (H) Comparison of whole genome of newly identified *K. weizmannii* to whole genomes of *Kazachstania* species

### In vitro characterization of *K. weizmannii*

When cultured in a number of defined conditions frequently found in the human body, such as serum exposure, neutral pH and / or 37°C, *C. albicans* forms hyphae ^12^. The same challenges did not invoke hyphenation in *K. weizmannii*, but the fungus continued to grow with yeast-like morphology (**Fig. 2A**).

**Figure 2:**
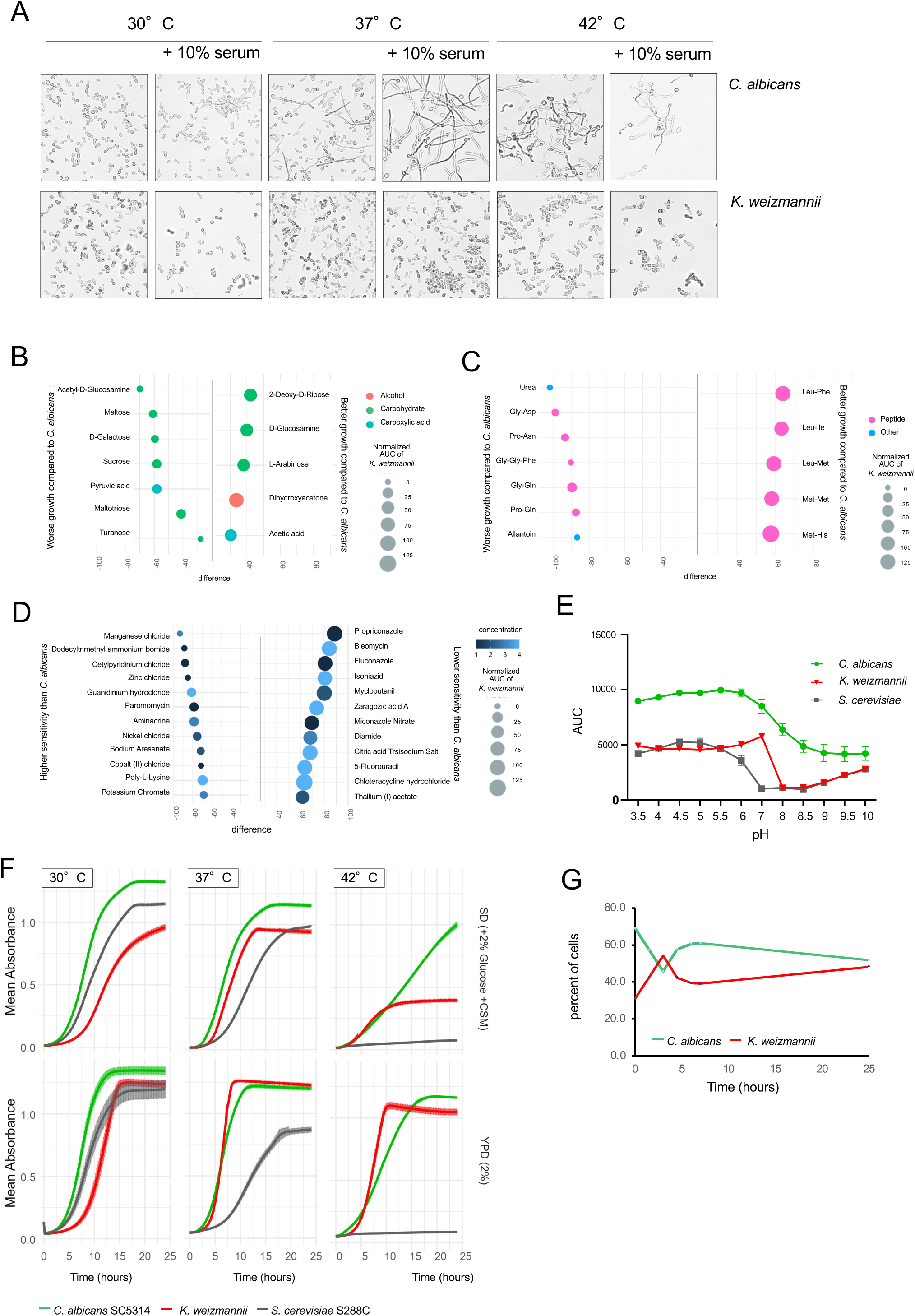
In vitro culture characteristics of *K. weizmannii*. (A) Representative images of *C. albicans* and *K. weizmannii* in liquid media in presence of filament-inducing conditions, n=3 independent experiments (B) Graph showing largest differences in growth between *K. weizmannii* and *C. albicans* with different carbon sources, based on a z-score > or < 2. Differences were calculated using the normalized area under the curve (AUC) of *K. weizmannii* and *C. albicans*. Normalization of the AUC based on the growth of both species with glucose. Size of data points corresponds to the normalized AUC of *K. weizmannii* with the corresponding carbon source. All measurements were done using the Biolog Phenotype MicroArrays in biological triplicates. X-axis represents the difference in AUC between *K. weizmannii* and *C. albicans*, while the y-axis shows the different carbon sources tested. The data points are color-coded according to the different types of carbon sources. (C) Graph showing largest differences in growth between *K. weizmannii* and *C. albicans* with different nitrogen sources, based on a z-score greater than or less than 2. Differences were calculated using AUC of *K. weizmannii* and *C. albicans*. Normalized based on the growth of both species with glutamine. Size of data points corresponds to the normalized AUC of *K. weizmannii* with the corresponding carbon source. All measurements were done using the Biolog Phenotype MicroArrays in biological triplicates. X-axis represents the difference in AUC between *K. weizmannii* and *C. albicans*, while the y-axis shows the different nitrogen sources tested. The data points are color-coded according to the different types of nitrogen sources. (D) Plot of the largest differences in growth of *K. weizmannii* and *C. albicans* with different inhibitors to test their chemical sensitivity. Graph shows the largest differences in growth between K. weizmannii and *C. albicans* with different inhibitors, based on a z-score greater than or less than 2., calculated using AUC of *K. weizmannii* and *C. albicans*. AUC normalization was based on the growth of both species without any inhibitor at pH 5. The size of each data point corresponds to the AUC of *K. weizmannii* with the corresponding inhibitor. All measurements were done using the Biolog Phenotype MicroArrays for chemical sensitivity (PM21-PM25) in biological triplicates. (E) Growth curves of *C. albicans, S. cerevisiae* and *K. weizmannii* under different pH conditions in YPD medium; AUC, Area under curve, biological triplicates (F) Growth curves of *C. albicans, S. cerevisiae* and *K. weizmannii* under different temperatures in SD medium (+2% Glucose+ CSM (complete supplement mixture)) and YPD, biological triplicates (G) Co-culture assay of *C. albicans* and *K. weizmannii* in liquid YPD medium. GFP expressing *C. albicans was* detected by flow cytometry to discriminate between fungi, n=2 independent experiments.

Comprehensive Biolog analysis (**Suppl. Data 1**) revealed differential preferences for carbon and nitrogen sources of *C. albicans* and *K. weizmannii* (**Fig. 2B, C**). *K. weizmannii* was, for instance, superior in growing with 2-Deoxy-D-Ribose and D-Glucosamine, as well as leucine dipeptides. Conversely, *C. albicans* thrived better on maltose, D-galactose, urea and glycine dipeptides. The two fungi also displayed differential resistance to chemical inhibitors (**Fig. 2D**). Specifically, *K. weizmannii* showed, as compared to *C. albicans,* relative resistance to propiconazole and fluconazole, while the yeast was more sensitive to chlorides and bromides. When cultured in SD media *C. albicans* and *K. weizmannii* displayed similar pH optima (**Fig. 2E**). Cultures under different temperatures revealed that both *K. weizmannii* and *C. albicans* grew at 37°C, while in line with it being a pathogen, *C. albicans* was superior in tolerating higher temperature (**Fig. 2F, Suppl. Fig. 2A**).

Interestingly and in contrast to the observation for the gut, *C. albicans* and *K. weizmannii* grew in *in vitro* YPD cocultures together without interference (**Fig. 2G, Suppl. Fig. 2B**). Under the condition tested competition between the strains was hence restricted to the gut environment.

### *K. weizmannii* is a murine commensal that antagonizes *C. albicans* colonization

To facilitate the comparative analysis of *K. weizmannii* and *C. albicans* SC5314, we generated a *K. weizmannii* strain harboring a gene encoding a red fluorescent miRFP reporter ^34^ in the enolase 1 (*ENO-1*) locus, mimicking the *C. albicans* SC5314-GFP configuration (**Fig. 3A Suppl. Fig. 3A,B**).

**Figure 3:**
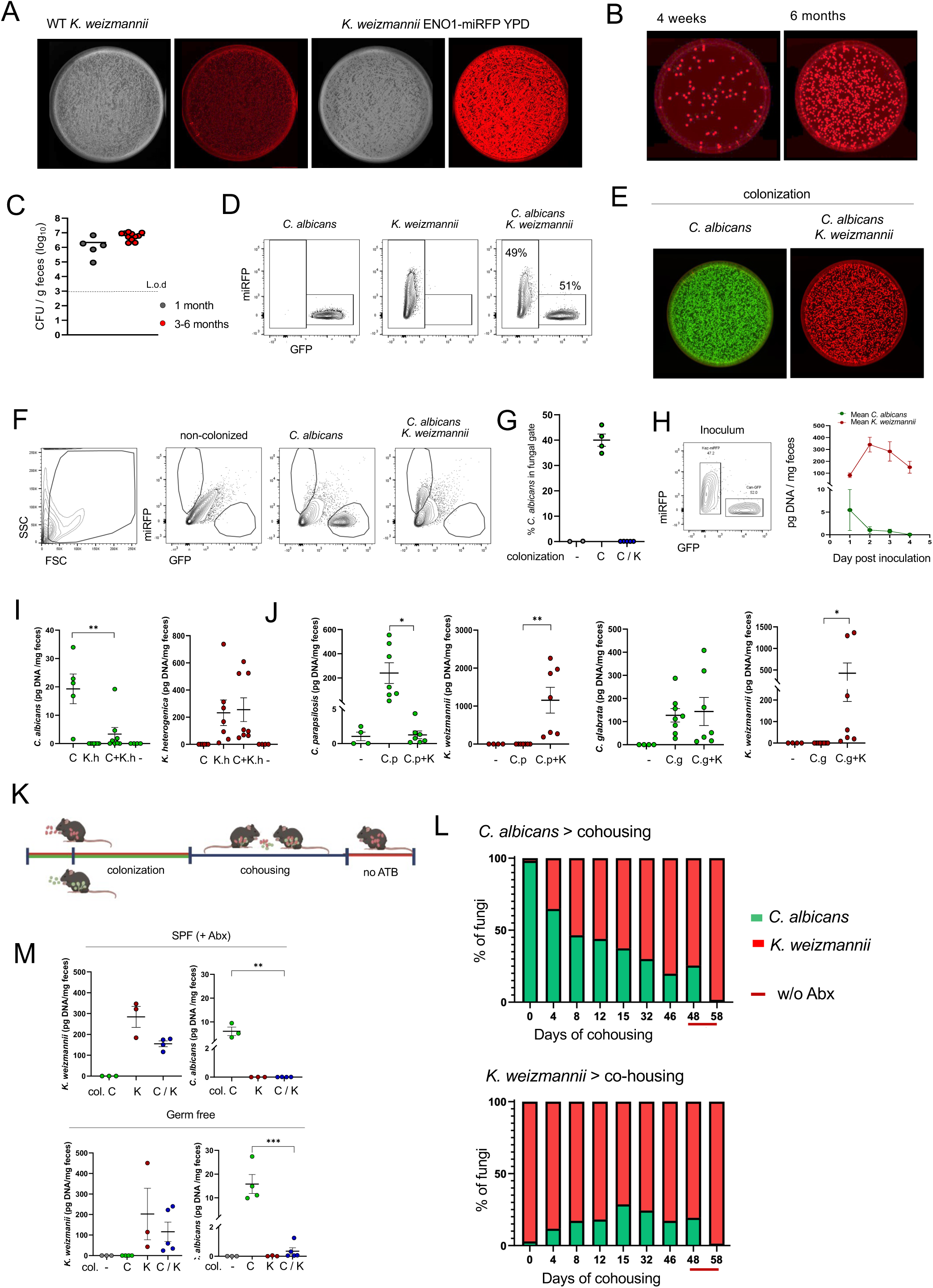
*K. weizmannii* competition with *C. albicans* in colonized animals. A) Far-red fluorescent reporter *Kazachstania* strain (*K. weizmannii* ENO1-miRFP). Representative picture. (B-C) Recoverable fecal *K. weizmannii* in colonized wild-type animals (kept without Abx) one and six months after single oral *K. weizmannii* ENO1-miRFP inoculation. (D) Flow cytometric analysis of *C. albicans* SC5314 (ENO1-GFP), *K. weizmannii* (ENO1-miRFP), and mixed cultures used for oral inoculation (E). (E) Representative plating analysis of feces of animals inoculated with 10^7^ yeasts of *C. albicans* SC5314 (ENO1-GFP) or *K. weizmannii* (ENO1-miRFP), and mixed cultures (see (D)) 3 weeks after administration. (F-G) Flow cytometric analysis of feces of animals orally inoculated with 10^7^ yeasts of *C. albicans* SC5314 (ENO1-GFP), *K. weizmannii* (ENO1-miRFP), and mixed cultures (see (D)) 3 weeks after administration. Quantification (n=3-5 mice per group, 4 independent experiments) (H) Flow cytometric analysis of a mixed culture used for oral inoculation (Right) and quantitative PCR analysis of genomic DNA of *C. albicans* and *K. weizmannii* recovered daily from feces of SPF animals colonized with a 1:1 mixture. Data shown as mean DNA quantities per mg of feces of N=5 Abx-treated mice, error bars represent SEM. (I) Quantitative PCR analysis of genomic DNA of *C. albicans* SC5314 and *K. heterogenica* recovered from feces of SPF animals colonized with either *C. albicans* SC5314 (C), *K. heterogenica* (K.h) or mixed culture (C+K.h) compared to Abx-treated SPF animals (-) 1 week after administration; 2 independent experiments. (J) Quantitative PCR analysis of genomic DNA of *C. parapsilosis, C. glabrata* and *K. weizmannii* recovered from feces of SPF animals colonized with either *C. parapsilosis* (C.p), *C. glabrata* (C.g) or mixed culture (C.p+K, C.g+K) compared to Abx-treated SPF animals (-) 1 week after administration; 2 independent experiments. (K) Schematic of co-housing experiment (L) Percentages of respective labelled fungi in feces of *C. albicans*, *K. weizmannii* colonized animals, before and after co-housing in Abx presence (except for day 48 and 58); Abx withdrawal on day 46 of cohousing, n=4 per group, representative of 2 independent experiments. (M) Quantitative PCR analysis of genomic DNA of *C. albicans* SC5314 and *K. weizmannii* recovered from feces of germ-free animals colonized with *C. albicans* SC5314 (C), *K. weizmannii* (K) or mixed culture (C/K) compared to Abx-treated SPF animals 1 week after administration; representative of 2 independent experiments.

Colonization of WT animals by *K. weizmannii* did not require prior Abx treatment, in contrast to the colonization with *C. albicans* in our facility **(Fig. 3B,C**). C57BL/6 mice could be readily colonized by oral *K. weizmannii* inoculation or cohousing with animals bearing the fungus. Interestingly, and in line with our original observation, when Abx-treated mice were co-inoculated with a 1:1 mixture of *K. weizmannii* and *C. albicans, Candida* colonization was prevented **(Fig. 3D-G**). Robust and rapid out-competition of *C. albicans* by *K. weizmannii* in this assay was confirmed by specific genomic PCR analysis of feces of respective co-inoculated animals (**Fig. 3H**). Competition was also observed with *K. heterogenica*, the yeast closest to *K. weizmannii* (**Fig. 3I**). Of note, competition was not restricted to *C. albicans*, but *K. weizmannii* co-inoculation also inhibited colonization with *C. parapsilosis*, an emerging fungal pathogen ^35^, but it did not interfere with *C. glabrata* colonization (**Fig. 3J**).

To test if *K. weizmannii* would also outcompete established commensal *C. albicans*, we stably colonized mice with *C. albicans* (on Abx) and then cohoused the animals with mice harboring *K. weizmannii*, maintaining Abx exposure **(Fig. 3K**). Coprophagy led to the progressive rapid ousting of *C. albicans* by *K. weizmannii*, leaving only a residual *C. albicans* population, which was further reduced upon Abx withdrawal (**Fig. 3L, Suppl. Fig. 3C**).

*C. albicans* colonization of mice is sensitive to the microbiome composition ^36,37^. The competition between the two fungi that we observed could hence be due to alterations of the bacterial landscape. Comparison of the microbiome of *K. weizmannii*-colonized mice and non-colonized controls using 16S sequencing revealed, however, only minor consistent changes (**Suppl. Fig 4A**). Moreover, also the abundance of Lactobacillae that are known to impede *C. albicans* colonization of the murine gut ^28^, was unaltered (**Suppl. Fig 4B**). To further probe the potential involvement of bacterial microbiota in the fungal competition, we orally inoculated germ-free animals with a mixture of the two fungi. Also in these mice, *K. weizmannii* prevented efficient *C. albicans* colonization (**Fig 3M, Suppl. Fig 4C-F**).

Collectively, these data establish that *K. weizmannii* can outcompete *C. albicans* including previously colonized animals and that the observed inter-fungal competition is independent of bacterial components.

### Comparative analysis of the host immune response to *C. albicans* and *K. weizmannii*

Fungi are known to affect host granulopoiesis ^38^ and induce both humoral and cellular immunity ^23,39^. In line with these reports, Abx-treated animals colonized with *C. albicans* showed an expanded blood neutrophil compartment. In contrast, *K. weizmannii*-colonized animals displayed no significantly altered abundance of granulocytes, classical or non-classical monocytes (**Fig. 4A**).

**Figure 4:**
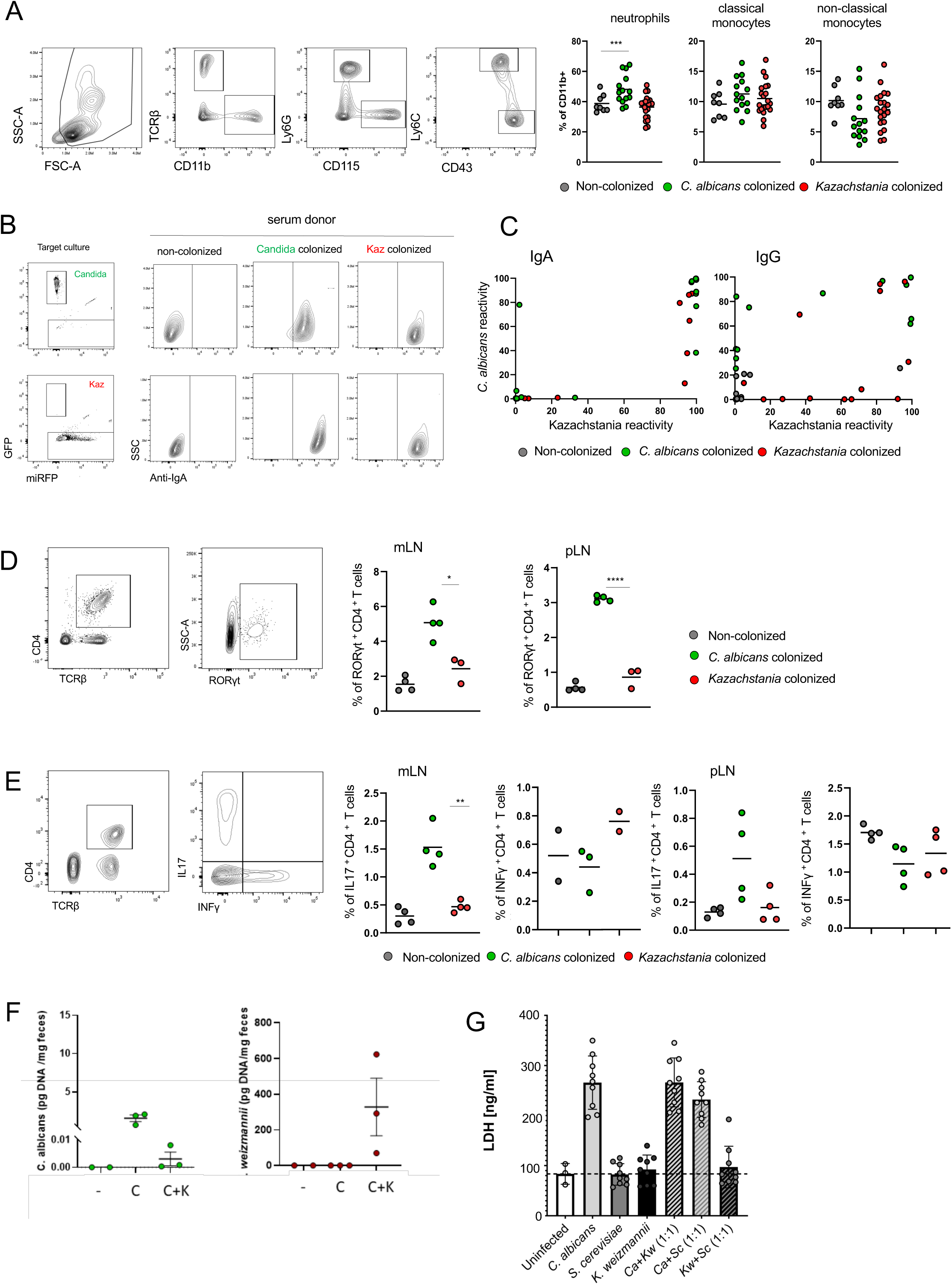
Humoral and cellular immune responses to *C. albicans* and *Kazachstania* sp. (A) Flow cytometric analysis of myeloid blood cell compartment of mice 4 weeks colonized with *C. albicans* or *K. weizmannii.* Neutrophils, classical and non-classical monocytes are defined as Ly6G^+^ CD115^-^, Ly6C^+^ CD115^+^, and Ly6C^-^ CD115^+^ cells, respectively. N= 8-21 mice, pooled from 3 independent experiments. Gating strategy, left, results, right. (B) Flow cytometric analysis of humoral anti-fungal reactivity. Gating strategy for the determination of serum immunoglobulin binding to cultured *C. albicans* or *K. weizmannii*. (C) Anti-*C. albicans* and *K. weizmannii* IgA and IgG serum reactivity of single-colonized animals, measured as shown in (B). Note that most but not all animals harbor cross-reactive sera. Data shown as percentage of Ig-positive gate, two pooled independent experiments, N=11-13 per group. (D) Representative gating strategy of RORγt**^+^** T cells (left); percentage of RORγt**^+^** cells among TCRβ CD4**^+^** T cells 1 month following *C. albicans* or *K. weizmannii* colonization, n=3-5 mice per group, 3 independent experiments (right) (E) Representative gating strategy of IL17 producing Th17 cells (left); percentage of IL17^+^ cells among TCRβ CD4**^+^**T cells 1 month following *C. albicans* or *K. weizmannii* colonization, n=3-5 mice per group, 3 independent experiments (right) (F) Quantitative PCR analysis of genomic DNA of *C. albicans* SC5314 and *K. weizmannii* recovered from feces of *Rag2^-/-^* animals colonized with *C. albicans* SC5314 (C), *K. weizmannii* (K) or mixed culture (C/K) 1 week after administration; representative of 2 independent experiments. (G) Epithelial cell (EC) coculture assay of *C. albicans*, *S. cerevisiae* and *K. weizmannii*. EC damage was assessed using the LDH assay as described in (Allert et al., 2018).

Intestinal IgA responses to *C. albicans* were proposed to balance commensalism vs. pathogenicity by controlling the critical morphological hyphae-to-yeast switch of the fungus^23^. To assess humoral immunity against the two fungal commensals, we analyzed sera of colonized animals for reactivity to cultured *K. weizmannii* or *C. albicans*. Colonization with either yeast induced robust anti-fungal serum IgA and IgG titers in most animals (**Fig. 4B, C, Suppl. Fig. 4G**). The induced antibodies were mostly, but not always, cross-reactive between the two fungal species, however, did not bind *S. cerevisiae* in the assay (**Suppl. Fig. 4H**).

Mucosa-associated fungi, and specifically *C. albicans* have been shown to induce Th17 type cellular immune response ^39,40^. Accordingly, *C. albicans*-colonized animals displayed an expansion of Th17 cells in gut mucosa-associated mesenteric and peripheral lymph nodes **(Fig. 4D, E).** In contrast, even after extended colonization, no Th17 cell expansion was observed in *K. weizmannii*-colonized animals. We did not observe significant alterations of the Th1 compartment in the LN analyzed with this assay, which however does not screen for antigen-specific cells **(Fig. 4E)**.

With the above we establish that colonized mice respond to the fungi, although the rapid kinetics of the out-competition we observe (**Fig. 3H**) suggest that the phenomenon is independent of adaptive cellular or humoral immunity. To directly test for potential role of B and T cells in the *Kazachstania / Candida* competition we investigated colonization in lymphocyte-deficient *Rag2^-/-^* mice. *K. weizmannii* prevented *C. albicans* colonization also in these lymphocyte-depleted animals (**Fig 4F**).

To directly gauge the impact of *K. weizmannii* on host cells we performed an epithelial cell (EC) co-culture assay (Allert et al., 2018). EC exposure to *C. albicans* resulted in EC damage as measured by LDH release. In contrast and similar to co-culture with *S. cerevisiae*, *K. weizmannii* did not affect EC viability **(Fig. 4G)**. However, co-infection of EC cultures with *K. weizmannii* and *C. albicans* did not prevent EC damage.

Taken together, colonization of Abx-treated animals with both *C. albicans* and *K. weizmannii* induced largely cross-reactive humoral immune responses reflected by serum IgA and IgG titers. The characteristic anti-fungal Th17 response was restricted to *C. albicans*-colonized mice, however the fungal competition we observe was independent of adaptive immunity. In combination with the results obtained from *in vitro* EC co-cultures, these data furthermore suggest that *K. weizmannii* is in mice an innocuous commensal.

### Commensal *C. albicans*, but not *K. weizmannii* causes pathology in immunosuppressed animals

Invasive candidiasis is widely recognized as a major cause of morbidity and mortality in the healthcare environment, often associated with an underlying immunocompromised state ^11,41^. Candidiasis can be induced in otherwise resistant, orally *C. albicans-*challenged animals by immunosuppression ^42^. To test whether corticosteroid treatment would cause *C. albicans* and *K. weizmannii* to spread from established commensal reservoirs and cause systemic pathology, we treated mice that were stably colonized with the respective fungi with cortisone 21-acetate boli (225 mg/kg s.c) every other day (**Fig. 5A**). Unlike non-colonized control mice, *C. albicans*-harboring animals lost significant weight after one week of treatment and became moribund (**Fig. 5B**). Animals displayed prominent tongue candidiasis and fungal growth in the kidneys (**Fig. 5C, D**). In stark contrast, immunosuppressed animals colonized with *K. weizmannii* showed neither weight loss nor evidence of fungal spread (**Fig. 5E-I, Suppl. Fig. 5A, B**). These data corroborate reports from the clinic ^11^ and mouse models ^43^ that gut commensal *C. albicans* is a pathobiont and can be the source of candidiasis. Furthermore, they establish that in mice, *K. weizmannii* is innocuous and even in immunosuppressed animals neither breaches the intestinal barrier to spread systemically nor causes other pathologies.

**Figure 5:**
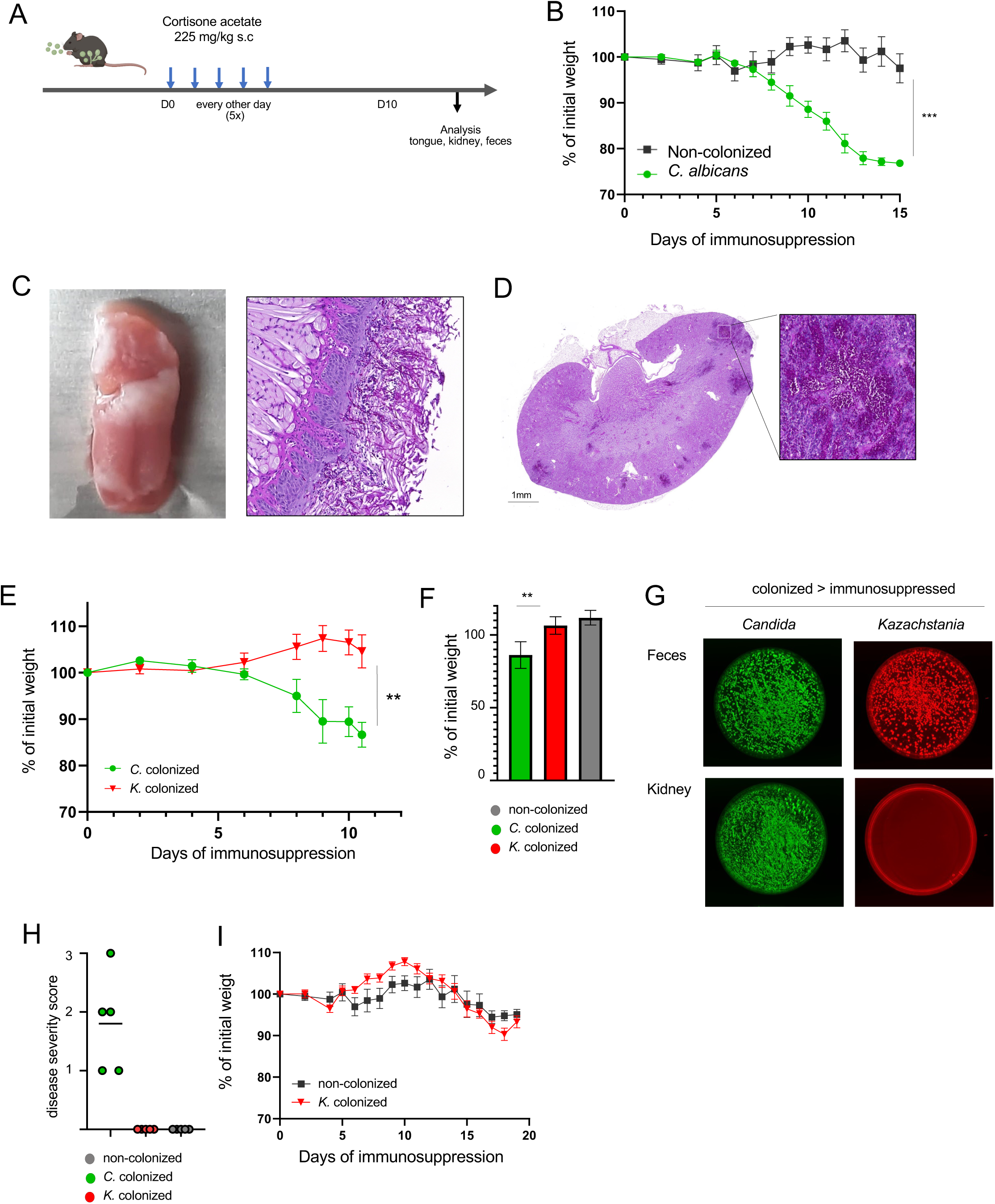
Commensal *C. albicans* but not *K. weizmannii* cause pathology in immunosuppressed animals. (A) Schematic of experimental setup for immunosuppression protocol. (B) Weight curve of *C. albicans*-colonized mice and uncolonized controls upon cortisone 21-acetate injections, n=5 mice per group, 3 independent experiments, error bar represents SEM. (C) Representative pictures of tongue candidiasis observed in *C. albicans*-colonized mice day 10 post immunosuppression, PAS staining (D) Representative picture of *C. albicans*-induced kidney pathology, PAS staining (E) Weight loss curve, comparison of *C. albicans* and *K. weizmannii* colonized mice, n=4-8 mice per group, 4 independent experiments, error bar represents SEM (F) Percent of initial weight in the endpoint of the experiment, graph represents mean with SEM (G) Representative picture of *C. albicans* and *K. weizmannii* recoverable by plating of feces (10 ng) and kidneys (10 mg) of *C. albicans* or *K. weizmannii* colonized animals (with Abx) 10 days after immunosuppression. (H) Pathology score based on microscopic and macroscopic evaluation of kidneys. (I) Weight loss comparison between *K. weizmannii*- and non-colonized animals. n= 5 per group, 3 independent experiments, error bar represents SEM

### Competitive commensalism mitigates Candidiasis

Since *K. weizmannii* exposure during competitive seeding and cohousing significantly reduced the commensal *C. albicans* burden in colonized animals (**Fig. 3J**), we next asked whether this commensal competition could mitigate candidiasis pathology. To simplify the mode of *Kazachstania* administration, we treated *Candida-*colonized animals with *Kazachstania*-supplemented drinking water **(Fig. 6A)** prior to immunosuppression. As expected, also in this setting *K. weizmannii* efficiently expelled *C. albicans* from colonized animals (**Fig. 6B, C**). Exposure of immunosuppressed *C. albicans*-colonized animals to *K. weizmannii* significantly delayed the weight loss and systemic yeast spread, as indicated by the absence of kidney colonization **(Fig. 6C-F, Suppl Fig. 6A)**. The animals were, though not permanently protected, as the residual commensal *C. albicans* eventually disseminated and caused pathology **(Fig. 6G, Suppl Fig. 6B)**.

**Figure 6:**
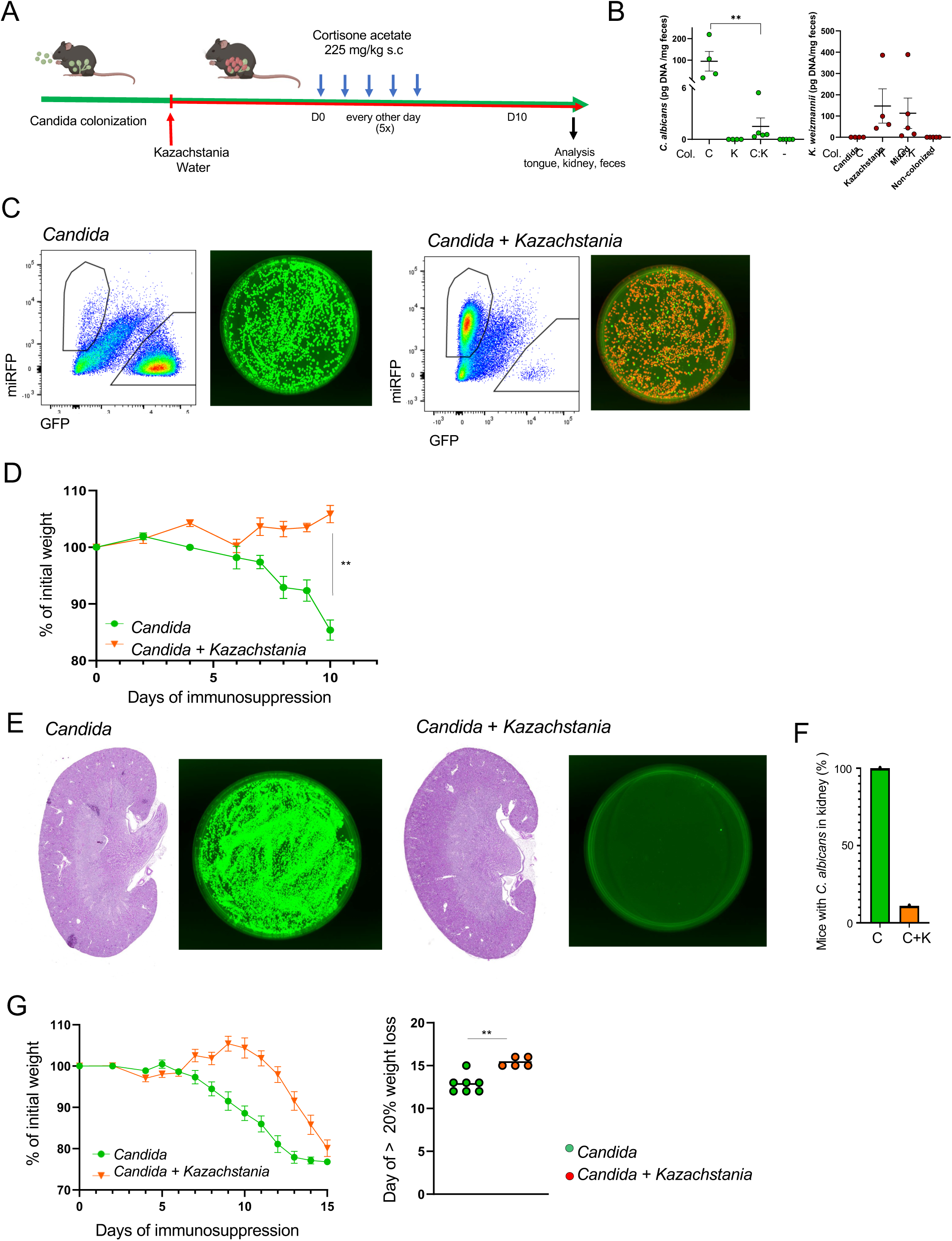
*K. weizmannii* mitigates *C. albicans*-induced pathology in immunosuppressed mice. (A) Schematic of immunosuppression protocol of *C. albicans*-colonized mice and oral *K. weizmannii* exposure. (B) Quantitative PCR analysis of genomic DNA of *C. albicans* and *K. weizmannii,* analyzed following 10 days of immunosuppression. Data shown as mean ± SEM, N=4-5 per group. (C) Representative picture of recoverable *C. albicans* and *K. weizmannii* from feces of *C. albicans*-colonized animals with or without *K. weizmannii* supplementation in their drinking water, analyzed by flow cytometry and cultivation 10 days after the first Cortisone-acetate injection. (D) Weight monitoring curve (% initial weight), comparison of immunosuppressed animals following *C. albicans*-colonization and *K. weizmannii*-outcompeted *C. albicans* colonization. All mice we sacrificed 10 days after the first dose of Cortisone-acetate injection. (E) Representative picture of kidneys (PAS staining) of immunosuppressed animals following *C. albicans*-colonization and *K. weizmannii*-outcompeted *C. albicans* colonization and results of cultivations of 1 mg kidney homogenate, kidneys were analyzed 10 days after the first Cortisone-acetate injection. (F) Bar graph indicating percentage of mice displaying C. albicans dissemination to kidneys as detected by plating and conformed by histology. *C. albicans* and *C. albicans* / *K. weizmannii*-outcompeted animals (N=9 per group). (G) Weight monitoring curve (% initial weight) – *C. albicans* and *K. weizmannii*-outcompeted animals, error bar SEM, comparison of the day reaching >20% weight loss

Collectively these results establish that *K. weizmannii* out-competes *C. albicans* from the commensal microbiome and by reducing the cause of pathology, mitigates candidiasis and improves the health status of immunosuppressed animals.

### *Kazachstania* presence in human microbiomes

*Candida* species are a major component of the human mycobiota, with *C. albicans* being most prevalent ^44^. Despite their established role in dough fermentation ^33^, *Kazachstania spp.* presence in healthy humans or during pathology-associated dysbiosis has rarely been reported ^45,46^. Accordingly, a recent study of human mucosa-associated mycobiota showed that these fungi are sparse among human commensals ^39^. To specifically gauge the abundance of *Kazachstania* spp., and in particular, our newly identified *K. weizmannii* in human microbiota, we designed a bioinformatic screen to detect genomic regions specific to these fungi, out of shotgun sequence information from a collection of 13,174 published metagenomics data sets ^2^ (**Suppl. Fig. 7A**). Among 7,059 human gut metagenome samples analyzed, we identified a few hundred samples that harbored ITS sequences of either the genus *Kazachstania* (i.e., with 9,236 read assemblies of 25s rRNA) or *Candida* (7,167 reads), or both (**Fig. 7A**). Among these, 32 metagenomes displayed specific evidence for *K. weizmannii* (20 unique to *K. weizmannii* and 12 shared with both species, listed in **Suppl. Fig. 7A**). Likewise, among 173 vaginal metagenomes analyzed, we found 22 samples with evidence for *Kazachstania* spp. and *Candida* spp. sequences, including 5 indicating specific presence of *K. weizmannii* (**Fig. 7B, Suppl. Fig. 7A**). *K. weizmannii-*positive samples had a wide range of biogeographical distribution (**Suppl. Fig. 7B, C**). Collectively, our data support the notion that *Kazachstania* spp. can be part of the human microbiome.

**Figure 7:**
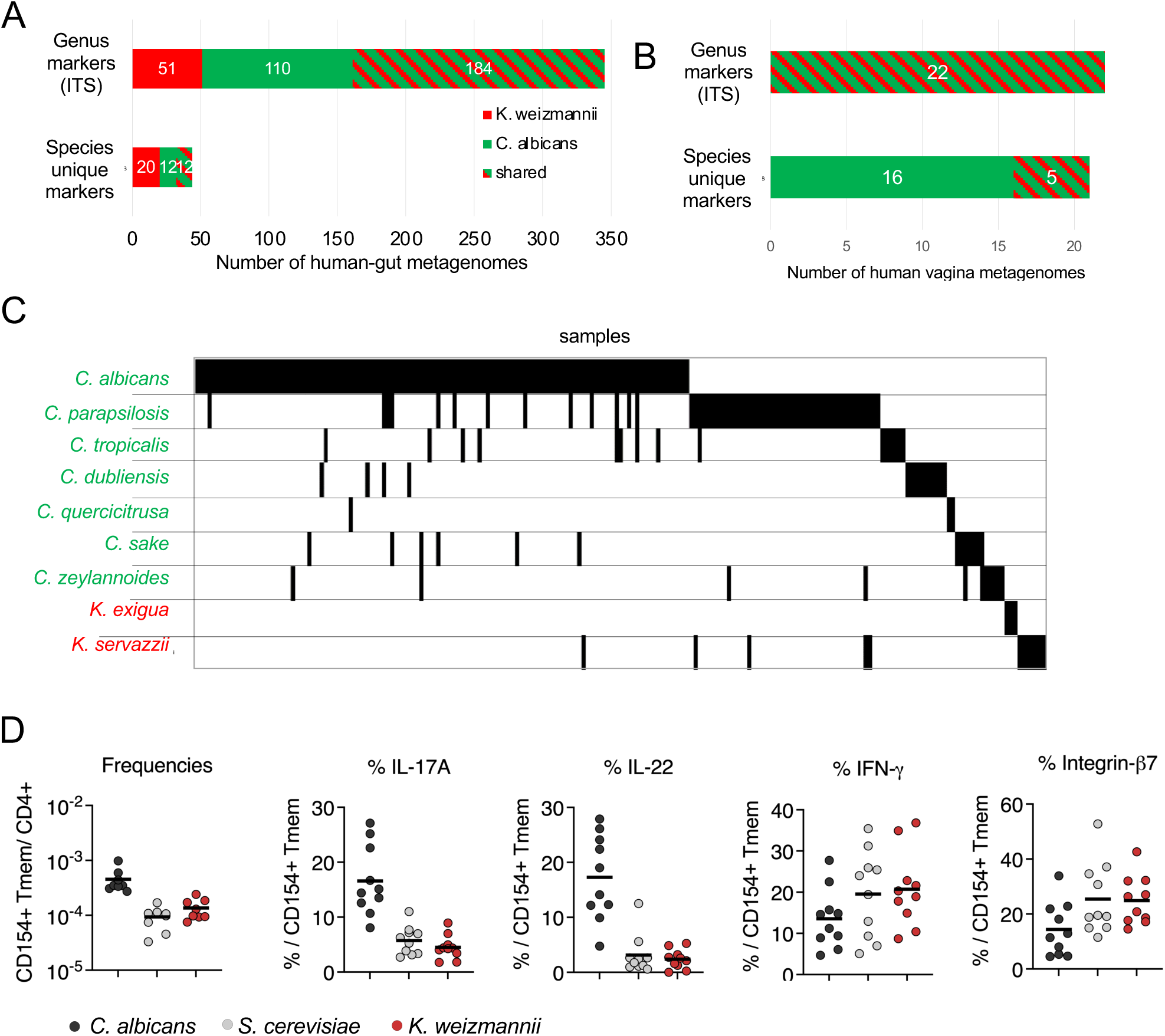
*Kazachstania* presence in human metagenomes. (A) Number of human gut metagenomes which were identified for either *C. albicans* (green) or *K. weizmannii* (red) or both (mixed color), using Genus-level markers (top) or using unique species-level markers (bottom). Numbers are out of 7059 examined metagenomic datasets. (B) Number of human vagina metagenomes which were identified for either *C. albicans* (green) or *K. weizmannii* (red) or both (mixed color), using Genus-level markers (top) or using unique species-level markers (bottom). Numbers are out of 173 metagenomic datasets. (C) ITS2 analysis of fecal samples of a cohort of 570 healthy individuals for presence of *Candida* and *Kazachstania* spp. (D) ARTE analysis of anti-fungal CD4^+^ T cell reactivity in blood of human individuals. (n= 10)

To further investigate the relative distribution of *Candida* and *Kazachstania* species in the human microbiome, we performed a sensitive ITS2 analysis of fecal samples of a cohort of 570 healthy individuals ^47,48^ (**Fig. 7C**). Since the *Kazachstania* genus had not been widely detected in prior ITS2 sequencing initiatives ^44^ we anticipated that it would be a less prevalent genus with lower abundance when compared with *Candida* and other prominent genera. Therefore, we added empty control samples which underwent the same amplification and library preparation processes to enable the application of a novel ITS2 processing pipeline previously applied to analyze the low biomass environment of the tumor mycobiome ^49^. Indeed, 13 fungal species could be detected in control samples with read numbers ranging from 1-100 reads per species per sample. We, therefore, applied an aggressive cutoff by flooring all species which obtained less than 100 reads in a given sample to zero in that sample. This process yielded 215 samples with ITS2 sequence evidence of either *Candida* or *Kazachstania* species. *C. albicans*, was the most prominent Candida species found in 120 samples, followed by *C. parapsilosis* and *C. tropicalis,* with 29 and 15 samples respectively. Two *Kazachstania* species were detected across 37 samples prior to flooring, and were completely absent from negative control samples. Still, these species were floored in samples where they did not reach 100 reads leaving 12 samples with *Kazachstania servazii* and 3 samples with *Kazachstania exigua,* close relatives of the novel *K. weizmannii* species. We did not detect *K. weizmannii* specifically in this limited cohort.

Finally, we corroborated *Kazachstania* presence in the human microbiome by Antigen-reactive T cell enrichment (ARTE) analysis ^50^ of peripheral blood of a limited number of healthy individuals. ARTE revealed T cell reactivity directed against *C. albicans* extracts, as shown earlier ^51^, but also against *K. weizmannii* (**Fig. 7D**). *C. albicans*-reactive T cells were polarized towards IL-17- and IL22-producing Th17 fates. In contrast, *K. weizmannii* - reactive T cells were INFγ-producing Th1 type cells, like T cells which reacted to *S. cerevisiae*. Of note, CD154^+^ memory T cells responsive to Saccharomycetaceae extracts also expressed β7 integrin indicative of their generation in the gut mucosa ^52^.

Collectively, these data establish Kazachstania spp. and specifically *K. weizmannii* as part of the human commensal microbiome.

## Discussion

Here we report the identification of a new fungal commensal of mice and human intestines. Unlike the well-studied *C. albicans, K. weizmannii* readily colonized animals kept under SPF conditions without prior Abx conditioning. Moreover, *K. weizmannii* efficiently outcompeted *C. albicans* when co-inoculated and also reduced the intestinal *C. albicans* load of previously colonized animals. Furthermore, competitive commensalism mitigated candidiasis pathology in mice under immune suppression.

Commensal mycobiota are understudied and rarely appreciated as a critical, active part of the microbiota. Noteworthy exceptions include the emerging evidence for the role of *C. albicans* in shaping Th17 immunity, behavior and potentially cancer progression ^39,49,53^. One of the reasons that commensal fungi have gained less attention than prokaryotes is arguably the fact that unlike mice roaming in the wild (Rosshart et al., 2017), animals kept under hyper-hygienic SPF conditions harbor poorly developed commensal mycobiota. Moreover, the study of fungi has primarily focused on *C. albicans*, but colonization of laboratory animals with this human pathobiont is impeded by bacterial communities, such as Lactobacillae ^28,29^ and therefore requires Abx conditioning ^27^.

*K. weizmannii* colonizes SPF mice without prior Abx conditioning and is hence, unlike *C. albicans,* resistant to bacterial commensals that impede fungal growth in SPF colonies ^28,29^. The mechanisms that underlie the competition of Lactobacillae with *C. albicans* are manifold and include toxic metabolic products, biofilm interference, and physical interactions and forced metabolic adaptation that compromises pathogenicity of *C. albicans* ^28,36^. Likewise, the mechanism that underlies competitive commensalism between *K. weizmannii* and *C. albicans* in the mice might be complex. The Biolog analysis for *in vitro* growth requirements revealed preferences of the respective fungi for specific carbon and nitrogen sources. Carbon and nitrogen catabolite metabolism has been discussed in the context of virulence of human pathogenic fungi ^54^. Whether the growth condition differences between *K. weizmannii* and *C. albicans* translate to the competition of the two fungi in the murine gut requires further study.

The results of the 16S analysis suggest that *K. weizmannii* does not significantly alter the composition of the bacterial microbiome of the host. However, the fungus might impact the fecal metabolomic landscape ^55^ by affecting bacterial expression profiles. Gastric *Candida* colonization of mice was shown to be affected by diet ^56^. Moreover, the fungal microbiome, including *Kazachstania* spp., was recently reported to be modulated by dietary intervention in pigs ^57^. The identification of *K. weizmannii* as a murine fungal commensal that colonizes mice in presence of bacteria should help to explore the impact of fecal procaryotic metabolome on mycobiota, their interactions, as well as the potential to manage colonization with pathobionts.

When co-housed with *K. weizmannii* bearing animals, mice under Abx treatment that are colonized with *C. albicans* progressively lose the pathobiont from the fecal microbiome. The residual *C. albicans* population is even further reduced upon Abx withdrawal. This finding might suggest the existence of two separate compartments in the intestine of mice: a fecal niche in which the fungi compete and an epithelia-associated niche which in SPF mice is dominated by commensal bacteria that prevent colonization by *C. albicans*, but not by *K. weizmannii*.

Following immunosuppression of colonized animals, *C. albicans* breaches the intestinal barrier ^58^, an activity which requires filamentation of the fungus ^59^, as well as expression of the cytolytic peptide toxin candidalysin that promotes penetration of the epithelial cell (EC) layer ^14^. In contrast, *K. weizmannii* remains confined to the gut lumen, in line with the observation that *K. weizmannii* do not form filaments upon in *vitro* stress and the fungus lacks part of the Core Filamentation Response Network identified in *C. albicans* ^60^.

We identified *K. weizmannii* by serendipity in mice that harbor impaired Th17 immunity; in line with its ability to colonize wt SPF mice without Abx conditioning, *K. weizmannii* was however widely spread in our animal facility irrespective of the genotype of the animals. Moreover, our metagenome screen yielded evidence that *K. weizmannii* is also part of human intestinal and vaginal microbiomes. Reports of *Kazachstania* spp. are however largely related to its involvement in food fermentation ^33^, while associations with murine or human commensalism or dysbiosis remain rare ^39,46^. It remains therefore unclear why the fungus so far escaped the radar. Rarefaction curves indicate that a cohort of 200 human samples is sufficient to comprehensively detect the human mycobiome taxonomic content, and our improved analysis pipeline pinpoints *Kazachstania* spp. as members. Yet, we observe that many members of the stool mycobiome, including *Kazachstania* spp., are masked by *S. cerevisiae* abundance and other food-related fungi. Only the arduous following of a serendipitous finding and the focus on a particular genus, enabled our finding. Notably, the majority of individuals harboring *Kazachstania* species in the local cohort displayed mutual exclusive presence with *Candida* spp. suggesting competitive fungal commensalism, as in mice. However, the notion of inverse correlation of these fungi and Candida spp. warrants further studies on a larger scale, since the sparsity of *Kazachstania* spp. precludes statistical significance.

Taken together, we identified with *K. weizmannii* an innocuous fungal commensal in men and mice. By its virtue to successfully compete with *C. albicans* in the murine gut for to-be-defined niches, *K. weizmannii* lowered the pathobiont burden and mitigated candidiasis development in immunosuppressed animals. This competitive fungal commensalism could have potential therapeutic value for the management of *C. albicans*-mediated diseases.

## Author contributions

J. S.K. made the initial observation and conceived the project with S.J., C.D. performed the yeast quantifications and serum titer analysis, S. B.H. S. T. provided animals and helped with analysis. S. B.D. and L. F. performed the genome analysis, B. D., I. L., L. N.-H., O. A., D. Z., E. S., Y. P. and R. S. contributed the human microbiome analysis. P. B. performed human T cell analysis, M.P.J., S. B. and B. H. performed the Biolog analysis. G. J. helped with yeast culture, O. B. performed histology, H. D. and N. S. helped with the sentinel screen and germfree analysis, N. S. advised on yeast biology and helped generating the reporter strain, P.B., B. H., S. B, J. S.K. and SJ wrote the manuscript.

## Acknowledgements

We would like to thank J. Berman (TAU) for providing the recombinant *C. albicans* (SC5314) reporter strain, Travis W. Adkins, (NRRL collection) for prompt yeast delivery, J. Zahradnik and G. Schreiber for the plasmid encoding the modified miRFP670 fluorescent protein, M. Lotan-Pompan and A. Weinberger for help with the bioinformatic analysis, S. Schäuble and M. Mirhakkak for providing the R script for the Biolog data analysis, C. Bar. Natan for help with the germ-free animal experiment and L. P. Coelho for helping with access to the metagenomes. SJ was funded by the Israeli Science Foundation (ISF) (grant # 696/21) and by the BINA (Bridge, Innovate, Nurture, Advance) program of the Weizmann Institute. This research was generously supported by Morris Kahn Institute for Human Immunology. S.J. is the incumbent of the Henry H. Drake Professorial Chair of Immunology. MJ, SB and BH were funded by the DFG through the Cluster of Excellence “Balance of the Microverse”, DFG project number 390713860. PB was funded by the Cluster of Excellence EXC2167 “Precision Medicine in Chronic Inflammation”, Project ID 390884018.

## List of Supplementary Materials

Supplementary Data 1. Biolog data

Supplementary Table 1. Details on microbiome screen

## Materials and Methods Mice

All animals involved in this study, unless otherwise noted, were on C57BL/6 background and of adult age (6-12 weeks). Mutant *Il23a^Δ/Δ^* mice were generated by crossing *Il23a^fl/fl^* animals ^61^ to *Pgk^Cre^* mice ^62^. Unless indicated otherwise, animals were maintained in a specific-pathogen-free (SPF) facility with chow and water provided ad libitum. Experiments were performed using sex- and age-matched controls. Animals were handled according to protocols approved by the Weizmann Institute Animal Care Committee (IACUC) as per international guidelines.

### Microbes

The recombinant C. albicans SC5314 strain expressing ENO1-GFP fusion protein was obtained from J. Berman (Tel Aviv University, Israel) ^30^. *K. weizmannii* was first cultivated from feces of mice housed in the Weizmann Institute SPF facility. The fluorescent *K. weizmannii* strain with a fusion of ENO1 to the modified miRFP 670 (herein Kazachstania-miRFP) was generated using CRISPR/Cas9 targeted mutagenesis ^30^. Insertion in ENO1 locus was confirmed using genomic PCR with primers ENO F (CGGTCAAATCAAGACTGGTGCTC) and miRFP Nano R1 (GCTGTTGCTGTTGCTGTAAAAGA) or Mirf R Screen (CTACCATGGGAGTATTCTTCTTCACC). *C. albicans* strain were cultured on solid YPD media at 30°C for 24h-36h. *K. weizmannii* and Kazachstania-miRFP were cultured on solid YPD media at 37°C. *K. heterogenica* Y-27499 was obtained from the ARS Culture Collection (https://nrrl.ncaur.usda.gov/) and cultured on solid YPD media at 37°C for 24h. *C. parapsilosis* (ATCC 2001) and *C. glabrata* were cultured on solid YPD media at 30°C for 24h.

### Fungal gut colonization

To establish intestinal colonization with *C. albicans* and *K. weizmannii* the drinking water of mice was supplemented with ampicillin (1 mg/mL, Ampicillin sodium salt 5 G SIGMA cat. 9518) 2-3 days prior to oral fungal inoculation. Mice were maintained on Abx-supplemented drinking water throughout the whole experiment, unless stated otherwise. For oral inoculation, *C. albicans* and *K. weizmannii* were grown on solid YPD media 30°C, or 37°C respectively. Cultures were washed with PBS, and 10^7^ yeast cells in 30 µl PBS were administered dropwise into the mouths of mice. For inoculation of mixed cultures, a culture of 10^7^ cells of each species was used. Non-colonized mice kept on ampicillin-supplemented water were used as control.

### Immunosuppression

Stably fungal-colonized mice were 5x s.c injected with 225mg kg^−1^ with cortisone 21-acetate (Sigma Aldrich C3130), following the scheme of injection every other day as described previously (Solis and Filler, 2012). Mice were monitored for weight loss. Following sacrifice, organs and feces were collected for histological examination, flow cytometry and fungal cultivations). If the weight dropped to 20% of the starting weight, mice were sacrificed according to IACUC protocol.

### Determination of CFU/g tissue/feces

For enumerating the number of recoverable *C. albicans* and *K. weizmannii* colony forming units, individual fecal pellets or each tissue from mice was sterilely dissected, weighed, and homogenized in sterile DDW. Serial dilutions on the organ homogenate were spread onto YPD media plates and the number of individual colonies enumerated after incubation at 37°C for 24 hours. Plates were imaged by Biorad ChemiDoc MP Imaging system.

### Genomic PCR for Yeast identification and quantification

Adopting a reported protocol ^53^, 25 mg of feces or tissue were surgically resected including its content. Samples were treated with Proteinase K, then homogenized using Lysing Matrix C (MP biomedicals) with the Omni Bead Ruptor 24 (Omni international, inc). DNA was extracted using Quick-DNA plus kit (Zymo Research) according to manufacturer’s instructions .10ng of isolated DNA were used for quantitative PCR reaction using Fast SYBR green Master Mix (Thermo Fisher Scientific, cat 4385614). Reaction was performed on the Quantstudio 7 Flex Real-Time PCR system (Thermo Fisher Scientific) using the fast SYBR 10µl program. Fungal DNA content in the samples was calculated using standard curves with known DNA concentration from cultured fungi and normalized to tissue weight. For detection of *C. albicans* we used previously published primers ^63^. For identification of fungi in colonized animals, feces were plated on YPD, following DNA purification from single colony and ITS1 detection via PCR reaction. PCR products were sequenced using in house Sanger sequencing.

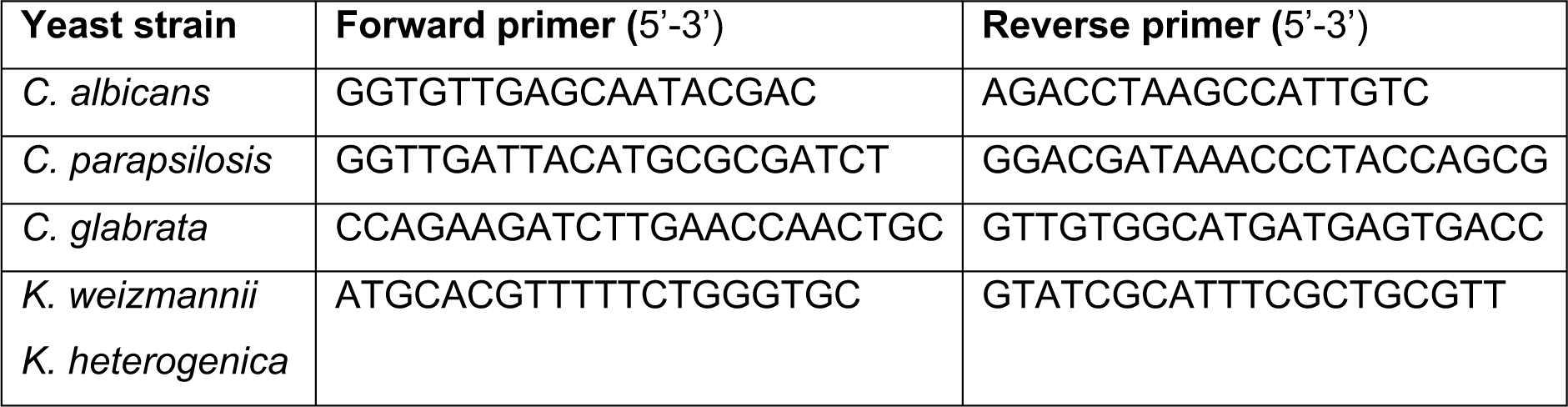

### Fecal DNA extraction for 16S rRNA gene sequencing

DNeasy Blood & Tissue Kit (cat. 69504) kit was used according to the manufacturer’s instructions for fecal DNA isolation for 16S sequencing. Prior the kit isolation, frozen fecal samples were digested with proteinase K in ALT buffer of the kit at 56°C, followed by bead-beating with sterile zirconia beads (Ø 0.1 mm, BioSpec, Cat. No. 11079101)

### 16S rRNA gene sequencing and taxonomic assignment

PCR amplification of fecal DNA was performed by Hylabs Company (Rehovot, Israel) using PCRbio Hot start ready mix using, using 2ul of DNA and custom primers covering the V4 region primers from Earth Microbiome Project (CS1_515F (V4_F), ACACTGACGACATGGTTCTACAGTGCCAGCMGCCGCGGT and CS2_806R (V4_R), TACGGTAGCAGAGACTTGGTCTGGACTACHVGGGTWTCT) for 25 cycles in a volume of 25ul. Of the reaction, 2ul was used for a second PCR amplification of 10 cycles in 10ul using Fluidigm Access Array Barcode library according to manufacturer’s protocol (2ul barcode per reaction). DNA was purified using Kapa Pure Beads at a ratio of 0.65X and quantified with Qubit fluorometer using Denovix DsDNA high sensitivity assay. DNA size and integrity was quantified by Agilent TapeStation DNA ScreenTape. Samples were sequenced on MiSeq (Illumina) machine with 30% PhiX using MiSeq Reagent Kit v2 500PE. Demultiplexing was performed using bcl2fastq with default parameters allowing for 0 mismatches. Data was then mapped to PhiX using bowtie2 to remove PhiX control and unmapped reads were quantified, collected and examined using fastQC. Demultiplexed reads were uploaded into CLC genomics workbench (Quiagen) and analyzed using their 16S microbiome pipeline. The analysis workflow consisting of quality filtration of the sequence data, and operational taxonomic unit (OTU) clustering was performed with default parameter settings. The adaptor sequence was removed and the reads with a quality score lower than 25 or length <150 were discarded. The maximum number of acceptable ambiguous nucleotides was set to 2 and the length of the reads was fixed at 200–500bp.

Chimeric sequences and singletons were detected and discarded. The remaining unique reads were used for OTU clustering, which was performed by alignment to the SILVA database at 97% sequence similarity.

### Bioinformatics analysis of 16S rRNA gene sequencing data

Visualization of OTU counts was done using the Marker Data profiling pipeline of MicrobiomeAnalyst ^64^. Counts were filtered to include OTUs with minimum 2 counts (mean abundance value) and scaled to library total sum. Abundance profiles were generated after merging small taxa with counts <10 based on their median counts. In order to explore uncultured species (D6 level), the annotations of ‘uncultured bacteria’ were concatenated to their family names (D4 level). Results were used for Alpha diversity analysis using Shannon diversity, T-test on filtered data, and Beta Diversity (PCoA using Bray-Curtis index) plots.

### Bioinformatics screening of human shotgun metagenomics datasets to identify fungi species

#### Identification of genomic regions unique to *K. weizmannii* and *C. albicans*

In order to identify *K. weizmannii* and *C. albicans* in human metagenomics datasets, we first selected a set of nucleotide regions that are unique to the genome sequence of either *K. weizmannii* or *C. albicans*, and were used to specifically identify these fungi in the background of other fungi, bacteria and other species in the metagenomes. The genome assemblies of *K. weizmannii* and *C. albicans* were compared to 22 genomes (genus Kazachstania) and 11 genomes (genus *Candida*), respectively (**Supplemental Tables 1-4**), using the GView Server ^65^, with analysis type ‘Unique genome’ with default pararmeters except for the Genetic code, where ‘Standard’ was used. Nucleotide regions unique to each target genome were collected, and further compared to the NCBI nt database using BLAST (E-value <0.0001) in order to exclude regions that are also present in bacteria or other non fungal organisms. This search resulted in 179 nucleotide sequences specific to *K. weizmannii*, and 904 nucleotide sequences specific to *C. albicans* (**Supplemental Table 1-4**).

#### Screening shotgun metagenomics datasets using unique *Kazachstania sp*. and *C. albicans* queries

A dataset of 13,174 metagenomes, collected from the Global Microbial Gene Catalog v1.0 (GMGC ^2^) was screened to identify sequences that originate from either *Kazachstania sp*. or *C. albicans* genomes. Each GMGC metagenome was originally stratified into habitats, and its raw nucleotide sequences were assembled into thousands of contigs (provided to us by Luis Pedro Coelho). All assembled contigs from each metagenomics sample were used as a query, and were searched against the set of unique *K. weizmannii* or *C. albicans* sequences constructed as described above, as a reference. Alignments were generated using bowtie2 algorithm (version 2.3.5.1, using --local mode). In order to broaden the search to genus level, all metagenomics assemblies were also mapped to a set of genus-specific rRNA and ITS sequences, extracted from *K. weizmannii* or *C. albicans* genomes (namely 25Sa, 18Sa, 5.8Sa, ITS1a, ITS2a, ETS1a, ETS2a). To avoid biases in the specificity of ITS searches, we counted only sequences which mapped to ITS regions without mutations (using the “XM:i:0” tag in the alignment file).

### Genomic DNA isolation from fungal cultures

Fungal pellet was dissolved in 2ml lysis buffer (100mM Tris pH 8.0, 50mM EDTA, 1% SDS) and sonicated for 10s. 300µl of the supernatant was transferred to 300µl 7M ammonium acetate pH 7.0, vortexed and incubated 5min at 65°C followed by 3 minutes on ice. 500µl chloroform was added and the solution was mixed by inverting the tube. Samples were spun 10min at 13000rpm at 4°C. 300µl of the upper phase was transferred to 400ul isopropanol-filled tubes and incubated for 5min on ice. Samples were spun 10min at 13000rpm at 4°C and pellet was washed with 800µl 70% EtOH, spun 1min at maximum speed, air dried and dissolved in DDW.

### Genome sequencing, annotation and comparison

Sequencing and hybrid (Nanopore and Illumina) assembly were performed by SeqCenter (Pittsburgh, PA), as follows: Illumina - Sample libraries were prepared using an Illumina DNA Prep kit and IDT 10bp UDI indices, and sequenced on an Illumina NextSeq 2000 producing 2−151bp reads. The data was demultiplexed and adapters removed using bcl2fastq [2] (v2.20.0.445) (https://support.illumina.com/sequencing/sequencing_software/bcl2fastq-conversion-software.html). Nanopore - Samples were prepared for sequencing using Oxford Nanopore’s “Genomic DNA by Ligation” kit (SQK-LSK109) and protocol. All samples were run on Nanopore R9 flow cells (R9.4.1) on a MinION. Basecalling was performed with Guppy (version 4.2.2), in high-accuracy mode (Default parameters +effbaf8). Quality control and adapter trimming was performed with porechop (https://github.com/rrwick/Porechop) version 0.2.2_seqan2.1.1 with the default parameters. Long read assembly with ONT reads was performed with flye (version 2.8) ^66^. The long read assembly was polished with pilon (1.23)^67^. Annotation was performed with the Yeast Gene Annotation Pipeline (YGAP) ^68^ (with the Post-WGD settings, and Companion for Fungi ^69^ with a reference organism of *Candida glabrata* CBS138. The output of both programs was compared with CD-HIT (version 4.8.1) with c=1, to reduce redundancy, and the remaining genes combined with YGAP as the base annotation, using in-house scripts. Whole genome comparison of 25 species of *Kazachstania* on the basis of *K. weizmannii* was performed with CCT (CGView Comparison Tool) ^66^, with a BlastN e-value of e-10.

### Phylogenetic Analysis

The d1d2 region of 26SrDNA of various species (**Supplemental Table 1**) were aligned with both ClustalW2.1 ^70^ and Muscle 3.8.31 ^71^. Phylogenetic trees were constructed with Maximum likelihood and DNA parsimony, using PhyML 3.0 ^72^, DNAML, DNAPARS and DNAPENNY in the Phylip package (3.697) developed by Felsenstein. The Phylip trees were then processed through Consense. Trees with similar topologies were obtained, and the Muscle/DNAPENNY tree is shown. The tree was visualized with iTol version 6 ^73^.

### In vitro serum antibody binding assay

Adopting a reported protocol ^23^, briefly, blood was collected from mice and rested at RT for 1 hour, followed by centrifugation 2000g 10 min. Serum supernatant was collected and froze at −80°C until use. Cultured fungi were normalized to OD_600_=1 in PBS supplemented with 1% bovine serum albumin and 0.01% sodium azide (PBA solution). Cultured fungi were incubated with 20x diluted mouse serum on ice for 45 minutes, then washed twice with PBA, followed by staining with anti-mouse IgA and IgG. Samples were recorded on Cytek Aurora and analyzed in FlowJo (Treestar). Antibody binding intensity was normalized to the negative controls. Isotype control, non-stained and no-serum were used as negative controls.

### Tissue isolation for flow cytometry

Mesenteric and peripheral lymph nodes were aseptically resected out into sterile, ice cold PBS and mashed manually using a 1 mL syringe plunger through a 80 µm nylon cell strainer. 100-200 µL of mouse blood was collected from submandibular vein, resuspended in 15 µl of heparin (Sigma) to prevent coagulation, followed by lysis with 1ml ACK buffer (8.29 g/ml NH4Cl, and 1 g/l KHCO3, 37.2 mg/l Na-EDTA) for 5 min RT and subsequent centrifugation.

### Cell staining, stimulation and flow cytometry and microscopy

Methods adhered to published guidelines ^74^. For cytokine staining,cells were incubated 3h with Cell Activation Cocktail (with Brefeldin A) (Biolegend, 423303) in10%FBS in RPMI at 37°C, followed by extracellular marker staining. Intracellular Fixation & Permeabilization Buffer Set (Invitrogen) was used according to manufacturer instruction. For transcription factor staining, Foxp3 /Transcription Factor Staining Buffer Set (Invitrogen) was used according to manufacturer instruction. Flow cytometry samples were recorded on BD LSR Fortessa 4 lasers or Cytek Aurora followed by data analysis using the FlowJo software (Treestar).

### Histology

Mice were euthanized and intestines, kidneys and tongue were excised and fixed overnight in 4% paraformaldehyde at 4°C. Paraffin embedding and sectioning was performed by the institutional histology unit. For histopathology of typical tongue or kidney fungal lesions in the mouse model of immunosuppression, paraffin sections were stained with periodic acid-Schiff (PAS) or H&E and slides were captured using a Panoramic SCAN II (3DHISTECH) and analyzed using CaseViewer software (3DHISTECH).

| REAGENT or RESOURCE | SOURCE | IDENTIFIER |
| --- | --- | --- |
| IgA, PE | Invitrogen | Cat#12-4204-81 |
| IgG, Alexa Fluor® 594 | BioLegend | Cat#405326 |
| CD45, Violet 450 | Biogems | Cat#07512-40-108 |
| CD4, Alexa Fluor® 647 | BioLegend | Cat#100412 |
| TCR $\beta$ chain, FITC | BioLegend | Cat#109206 |
| IgG1 $\kappa$ Isotype Ctrl, PE | BioLegend | Cat#400407 |
| ROR gamma (t) | Invitrogen | Cat#12-69981-80 |
| IFNγ, APC | BioLegend | Cat#505810 |
| IL17, PE | BioLegend | Cat#506904 |
| TCR $\beta$ chain, PE/Cyanine5 | BioLegend | Cat#109209 |
| CD274, PE/Dazzle™ 594 | BioLegend | Cat#124323 |
| CD43, Alexa Fluor® 700 | BioLegend | Cat#143214 |
| Ly6C, Pacific Blue™ | BioLegend | Cat#128014 |
| Ly6G, FITC | BioLegend | Cat#127605 |
| CD115, APC/Cyanine7 | BioLegend | Cat#135532 |
| CD11b, PE | BioLegend | Cat#101207 |
| CD220, APC | BioLegend | Cat#103212 |
| Zombie Aqua™ Fixable Viability Kit | BioLegend | Cat#423102 |
| Cy3-AffiniPure Donkey Anti-Rabbit | J. Immunoresearch | Cat#711165152 |
| purified CD16/32 | BioLegend | Cat#101302 |
| Cell Activation Cocktail | BioLegend | Cat#423302 |
| Brefeldin A Solution | BioLegend | Cat#420601 |
| Fixation/Permeabilization Kit | BD Bioscience | Cat# 554714 |
| True-Nuclear™ TF Buffer Set | BioLegend | Cat#424401 |

### Antigen-reactive T cell enrichment (ARTE)

PBMCs were freshly isolated from EDTA blood samples on the day of blood donation by density gradient centrifugation (Biocoll; Biochrom, Berlin, Germany). Antigen-reactive T cell enrichment (ARTE) was performed as previously described ^50,75^. In brief, 2×10e7 PBMCs were plated in RPMI-1640 medium (GIBCO), supplemented with 5% (v/v) human AB-serum (Sigma Aldrich, Schnelldorf, Germany) at a cell density of 1×10e7 PBMCs / 2 cm^2^ in cell culture plates and stimulated with 40µg/ml fungal lysates for 7 hr in presence of 1 µg/ml CD40 and 1 µg/ml CD28 pure antibody (both Miltenyi Biotec, Bergisch Gladbach, Germany). 1 µg/ml Brefeldin A (Sigma Aldrich) was added for the last 2 hr. Cells were labeled with CD154-Biotin followed by anti-Biotin MicroBeads (CD154 MicroBead Kit; Miltenyi Biotec) and magnetically enriched by two sequential MS columns (Miltenyi Biotec). Surface staining was performed on the first column, followed by fixation, permeabilization (Inside stain Kit, Miltenyi Biotec) and intracellular staining on the second column. The following antibodies were used: CD4-APC-Vio770 (M-T466), CD8-VioGreen (REA734), CD14-VioGreen (REA599), CD20-VioGreen (LT20), Integrin-b7-PE-Vio770 (REA441) (all Miltenyi Biotec); CD45RA-PE-Cy5 (HI100), IFN-γ-BV785 (clone: 4S.B3) (both Biolegend); IL-17A-BV650 (clone: N49-653), IL-22-PerCP-eFluor710 (clone: IL22JOP) (both BD Biosciences). Viobility 405/520 Fixable Dye (Miltenyi Biotec) was used to exclude dead cells. Data were acquired on a LSR Fortessa (BD Bioscience, San Jose, CA, USA).

Frequencies of antigen-specific T cells were determined based on the total cell count of CD154+ T cells after enrichment, normalized to the total number of CD4+ T cells applied on the column. For each stimulation, background cells enriched from the non-stimulated control were subtracted.

### ITS2 amplification and sequencing of human stool samples

ITS2 sequencing was used for fungal identification as described in ^49^. Briefly, ITS2 sequencing applied to 570 human stool samples and 6 controls ^48,76^. PCR was performed on 10ng of DNA per sample (or the maximum available). Three PCR batches were required with 2 wells left empty as library controls in each batch. Forward primer ITS86F 5’-759 GTGAATCATCGAATCTTTGAA-3’ and reverse primer ITS4 with rd2 Illumina adaptor 5’-AGACGTGTGCTCTTCCGATCT - TCCTCCGCTTATTGATATGC-3’ were used for the first PCR amplification. PCR mix per sample contained 5ul sample DNA, 0.2μM per primer (primers purchased from Sigma), 0.02 unit/μl of Phusion Hot Start II DNA Polymerase (Thermo Scientific 763 F549), 10μl of X5 Phusion HS HF buffer, 0.2mM dNTPs (Larova GmbH), 31.5ul ultra pure water, for a total reaction volume of 50μl. PCR conditions used were 98°C 2min, (98°C 10 sec, 55°C 15 sec, 72°C 35 sec) X 30, 72°C 5 min. A second PCR was performed to attach Illumina adaptors and barcode per sample for 6 additional cycles.

Samples from 1st PCR were diluted 10- fold and added to the PCR mix as described above. Primers of second PCR included: forward primer P5-rd1-768 ITS86F 5’ - AATGATACGGCGACCACCGAGATCTACACTCTTTCCCTACACGACGCTCTTCCGATCT-GTGAATCATCGAATCTTTGAA-3’, and reverse primer 5’- CAAGCAGAAGACGGCATACGAGAT - NNNNNNNN - GTGACTGGAGTTCAGACGTGTGCTCTTCCGATCT-3’. Every 96 samples were combined for a single mix by adding 14μl from each. Before mixing, an aliquot from each of the samples was run on an agarose gel. In cases where the amplified bands were very strong, samples were diluted between 5 and 20-fold before they were added to the mix. Each sample mix was cleaned with QIAquick PCR purification kit (QIAGEN, catalog # 28104). Two cleaned sample mixes were then combined into a single mix of 192 samples, and size selection was performed with Agencourt AMPure XP beads (Beckman Coulter #A63881) to remove any excess primers. Beads to sample ratio was 0.85 to 1. Samples were then run in three libraries on the Miseq v3 600 cycles paired-end with 30% PhiX.

### ITS2 sequencing analysis

The ITS2 classification pipeline was built with Python 3.6. For each sequencing library, paired-end reads were joined using PEAR (version 0.9.10) followed by filtering of merged reads by minimum length of 80bp and trimming of primers from both ends with cutadapt (version 1.17). Within the QIIME 2 environment (version 2018.8), Dada2 was used to create amplicon sequence variants (ASVs), then ITSx (version 1.1b1) was used to delineate ASVs to ITS2 regions (removing preceding 5.8S and trailing 28S sequences). A taxonomic naive bayesian classifier in QIIME 2 ^77^ was trained on the UNITE database (version 8, dynamic, sh_taxonomy_qiime_ver8_dynamic_04.02.2020.txt) and used to classify the 180 processed ASVs. 91 percent of raw reads were classified to species level (**Table 1**).

Most of the downstream analysis and plots were performed with R version 4.1.1 and phyloseq 1.34.0. ASVs were filtered by the ITSx and UNITE classifications to include fungal reads only. 119 ASVs that were classified by ITSx as fungi were included in the downstream analysis, representing over 97.5% of reads. Out of the remaining 61 ASVs that were classified by ITSx as non-fungal (Tracheophyta (T), land plants), one (fid65) was included in the downstream analysis since its classification as fungi reached all the way to species level by UNITE and was validated by NCBI BLAST to be fungal. The histogram of the number of reads per ASV per sample presented a bimodal distribution with the peaks found on either side of 100 reads/ASV or 100 reads/sample. We therefore floored the data in a sample-specific manner, such that if an ASV was assigned less than 100 reads in a specific sample, its assigned reads were converted to zero. Next, we introduced two types of data normalization: (1) Library normalization, where samples were normalized to account for the difference in the average number of reads/sample per library. (2) Dilution normalization: ASV reads were multiplied by the dilution factor per sample to reflect their true original load. Next, ASVs were aggregated based on UNITE classification, to species level when possible. ASVs that could not be classified to species level, were grouped together by the lowest known phylogenetic level and labeled “Other”. A total of 55 species were detected, 13 of which were also detected in control samples, but with a maximum of 100 reads per sample demonstrating a very mild read leakage of the higher abundant species in the library, e.g. Saccharomyces cerevisiae. Therefore, flooring of species with less than 100 reads per sample was applied. Lastly, data were aggregated by summing all reads in each taxonomic level by the associated taxa in the level above it.

**Table 1.**

| **Table 1** |  |  |
| --- | --- | --- |
|  | **number** | **percent** |
| **kingdom** | 29339412 | 100 |
| **phylum** | 29339412 | 100 |
| **class** | 29339412 | 100 |
| **order** | 29339412 | 100 |
| **family** | 28453602 | 96.98082 |
| **genus** | 28423083 | 96.8768 |
| **species** | 26935819 | 91.80763 |

### Strains and culturing conditions

*Candida albicans* SC5314, *Saccharomyces cerevisiae* S288C, and *Kazachstania weizmannii* were regularly cultured in YPD (1% yeast extract, 2% peptone, 2% glucose) medium or on agar plates at 30 °C or 37 °C. Precultures were prepared in liquid YPD and incubated at 30°C with shaking until stationary (approx. 22 h). For growth experiments, cultures were washed twice with PBS, and adjusted to an experiment-specific OD in PBS. Where indicated, synthetic defined medium was used (SD; 6.7 g Yeast Nitrogen base (YNB) + ammonium sulfate (Formedium, Great Britain), 20 g glucose, with or without 0.79 g complete supplement mixture (CSM) (Formedium, Great Britain) in 1 l of distilled water).

### Growth in different media and temperatures

Growth was measured in 96-well format in these media: YPD (1% yeast extract, 2% peptone, and 2% glucose), BHI (“brain heart infusion”, Roth), SD (Formedium, Great Britain) with 2% glucose ±CSM, DMEM (Dulbecco’s Modified Eagle Medium, Gibco, Thermo Scientific) or RPMI 1640 (Gibco, Thermo Scientific). The initial OD was set to 0.1 using 200 µl as the total volume of media per well. Cell-free medium served as control. Each experiment was done in three biological replicates per strain and per media. The 96-well-plates were incubated at either 30, 37, or 42°C and growth was measured optically at 600 nm in a BioTek LogPhase 600 multiplate reader (Agilent Technologies, Inc., USA) every 10 minutes for 24 hours. Mean growth of the replicates was visualized by R version 4.2.2 (R Core Team, 2022, https://www.R-project.org/).

### Virulence of *K. weizmannii*

Host cell damage was determined with C2BBe1 human intestinal epithelial cells (ATCC CRL-2102) by release of cytoplasmic LDH. The intestinal cells were seeded collagen I-coated (10 μg/ml, 2 h at room temperature [RT]; Thermo Scientific) 96-well plates at a total cell density of 1×10^5^ cells/ml in 200 µl DMEM, supplemented with 10% fetal bovine serum (FBS; Bio & Sell), 10 μg/ml Holotransferrin (Calbiochem, Merck), and 1% non-essential amino acids (Gibco, Thermo Scientific), then incubated for 48 hours at 37°C and 5% CO_2_. Overnight yeasts cultures were semi-synchronized by diluting in YPD to an OD of 0.2, followed by 4 h at 30°C, 180 rpm shaking. The precultures were then washed twice with 1 ml sterile PBS and dissolved in 1 ml DMEM, and adjusted to 8×10^5^ cells/ml in DMEM singly or, as coinfection, each. The medium was removed from the C2BBe1 cells by aspiration, 100 µl DMEM without FBS and 100 µl of the yeast suspensions (or DMEM alone) were added in technical triplicates. The cells were incubated for 24 hours at 37°C with 5% CO_2_. Host cell damage was detected with the Cytotoxicity Detection Kit (Roche) from the supernatants according to the manufacturer’s instructions and calibrated by LDH from rabbit muscle (5 mg/ml, Roche). Data analysis was done with GraphPad Prism.

### Phenotypic screening

The Phenotype MicroArrays for microbial cells (PM) system (Biolog Inc., USA) was used according to the manufacturer’s instructions. Colonies were transferred with sterile cotton swaps into 15 ml of sterile dH_2_O at a turbidimeter transmittance of 62%. PM plates used were carbon sources (PM1-2), nitrogen sources (PM3, PM6-8), pH (PM10), and chemical inhibitors (PM21-25). Extended Data 1-list all tested substances. For carbon and nitrogen sources, the medium contained inoculating fluid IFY-0 base, redox dye mix D (Biolog Inc., USA), potassium phosphate, and sodium sulfate, supplemented with L-glutamic acid monosodium (for PM1-2) or glucose (for PM3, PM6-8) (Sigma-Aldrich). For inhibitors and pH (PM10, PM21-25), SD medium (Formedium, Great Britain) supplemented with CSM (Formedium, Great Britain), redox dye mix E (Biolog Inc., USA), ammonium sulfate, and glucose was used. Metabolic activity of 100 µl yeasts suspension was followed as color reaction over 48 h at 37°C every 15 minutes (Mackie et al., 2014) in biological triplicates.

Data were processed using Biolog Data Abn alysis 1.7 (Biolog Inc.,USA) and analyzed by a previously described R pipeline ^78^ and the package opm version 1.3.77 ^79^ to group into active)log growth) and non-active, and calculate the AUC. AUCs were normalized for each species to growth with either glucose (for carbon sources), glutamine (nitrogen sources) or growth with inhibitor-free test medium at the same pH of 5 (inhibitors). The difference between the normalized AUCs of C. albicans and K. weizmannii

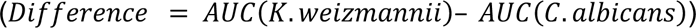 was used to calculate the z-scores 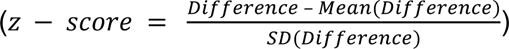. Substances with z-scores > 2 or < −2 were graphed with ggplot2 (https://ggplot2.tidyverse.org).

### Filamentation assay

*C. albicans* and *K. weizmannii* were tested for their ability to the filament in liquid filamentation conditions. Yeast cells grown overnight in liquid YPD at 30 °C or 37 °C respectively, were centrifuged and washed 2× with PBS prior to the experiment and strains were diluted on the same OD. Yeast cells were incubated for 5h in 30°C, 37°C or 42°C with shaking, then directly fixed with 4% PFA in PBS for 30 min, followed by wash with PBS and images were captured by bright Zeiss field microscope.

### Statistical analysis

In all experiments, data are presented as the mean ± SEM unless stated otherwise. Statistical significance was defined as P < 0.05. The number of animals is indicated as n. Animals of the same age, sex, and genetic background were randomly assigned to treatment groups.

### Data availability

The 16S sequencing data have been deposited to BioProject accession number PRJNA949686 in the NCBI BioProject database (https://www.ncbi.nlm.nih.gov/bioproject/)

---

**Suppl. Figure 1:**
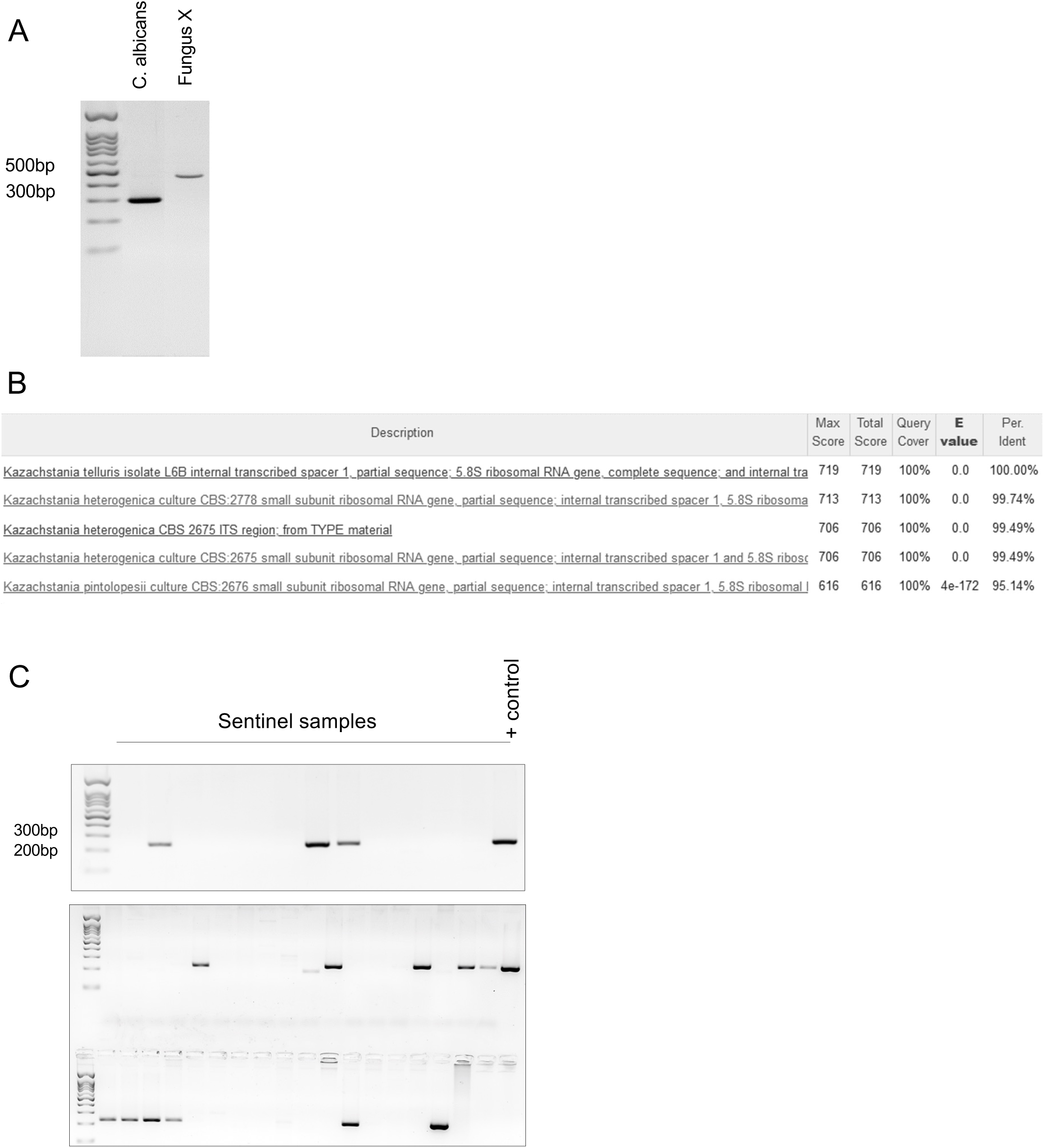
Identification of novel fungal isolate. (A) Gel analysis of PCR products amplifying ITS1 region of *C. albicans* and a fungal isolate (B) Identification of novel fungal sp. based on ITS1 sequence (C) Representative example of PCR sentinel screening using PCR primers for specific ITS1 of *K. weizmannii*.

**Suppl. Figure 2:**
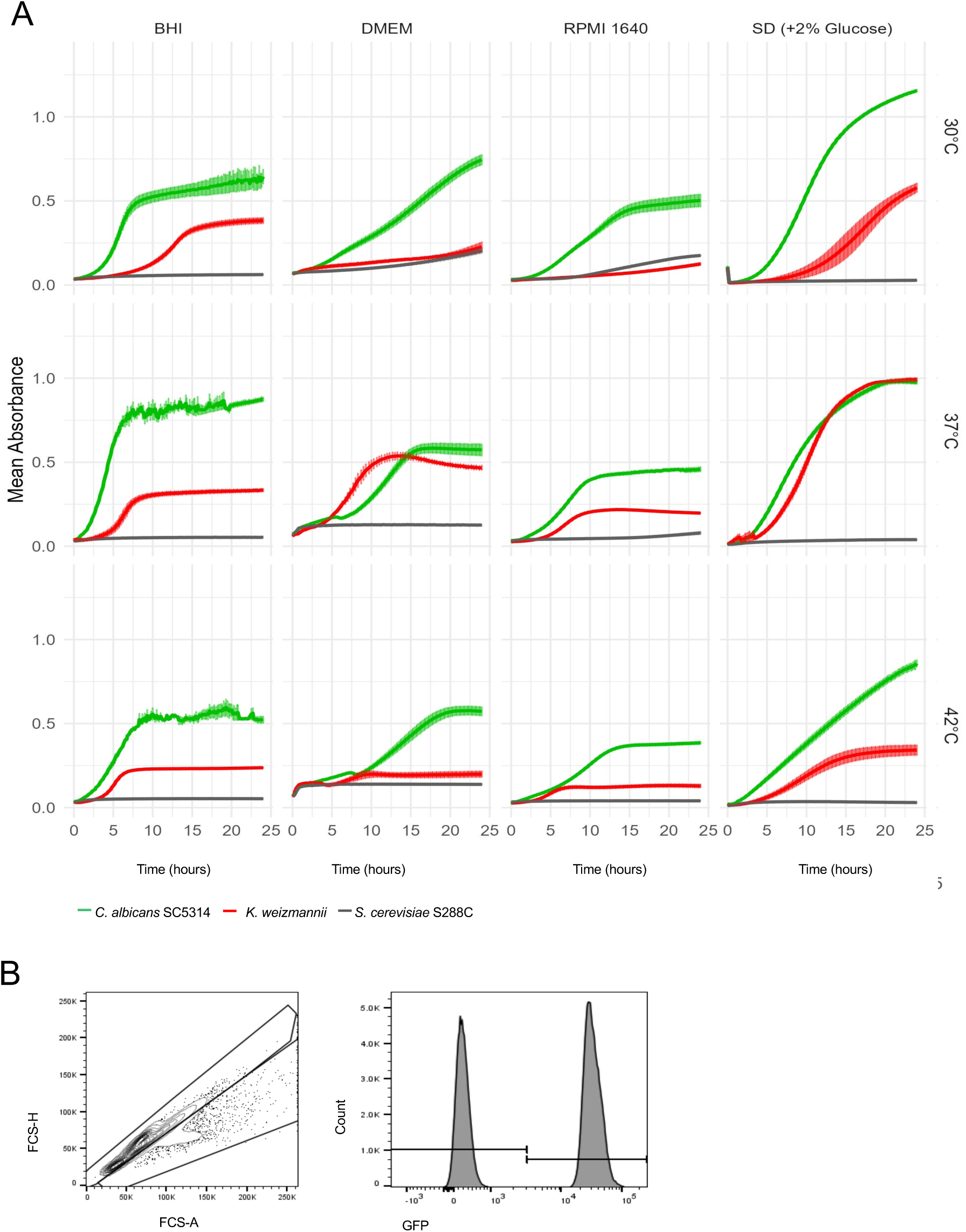
Generation of *K. weizmannii* reporter strain. (A) Temperature growth curves of *C. albicans, S. cerevisiae* and *K. weizmannii* under different temperatures in indicated media, biological triplicates (B) Example of flow cytometric analysis of mixed culture assay of *C. albicans* and *K. weizmannii* taking advantage of GFP expressing *C. albicans*

**Suppl. Figure 3:**
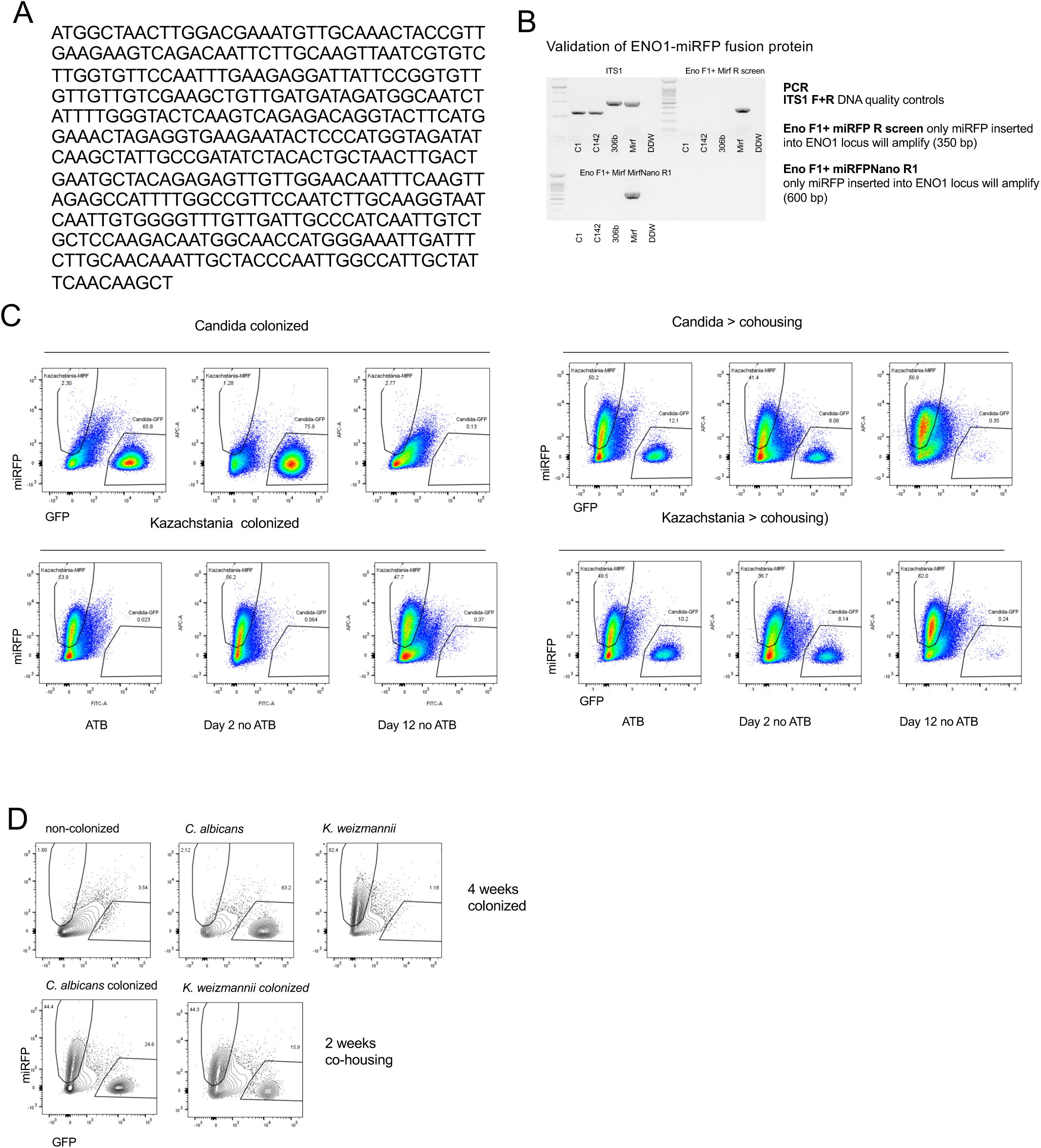
**Generation of *K. weizmannii* reporter strain**. (A) Sequence of modified fluorescent reporter miRFP670 – further referred as ‘miRFP’ (B) PCR validation of proper insertion of miRFP into *K. weizmannii* genome and thus generation of ENO1-miRFP fusion protein. (C) Representative picture of recovery of *C. albicans* ENO1-GFP and *K. weizmannii* ENO1-miRFP from feces of single-colonized animals followed by cohousing (started as Candida or Kazachstania) (D) Representative flow cytometric feces analysis of colonized mice 4 weeks after initial colonization (before cohousing) and 2 weeks after co-housing. (N=4 per group, 2 independent experiments)

**Suppl. Figure 4:**
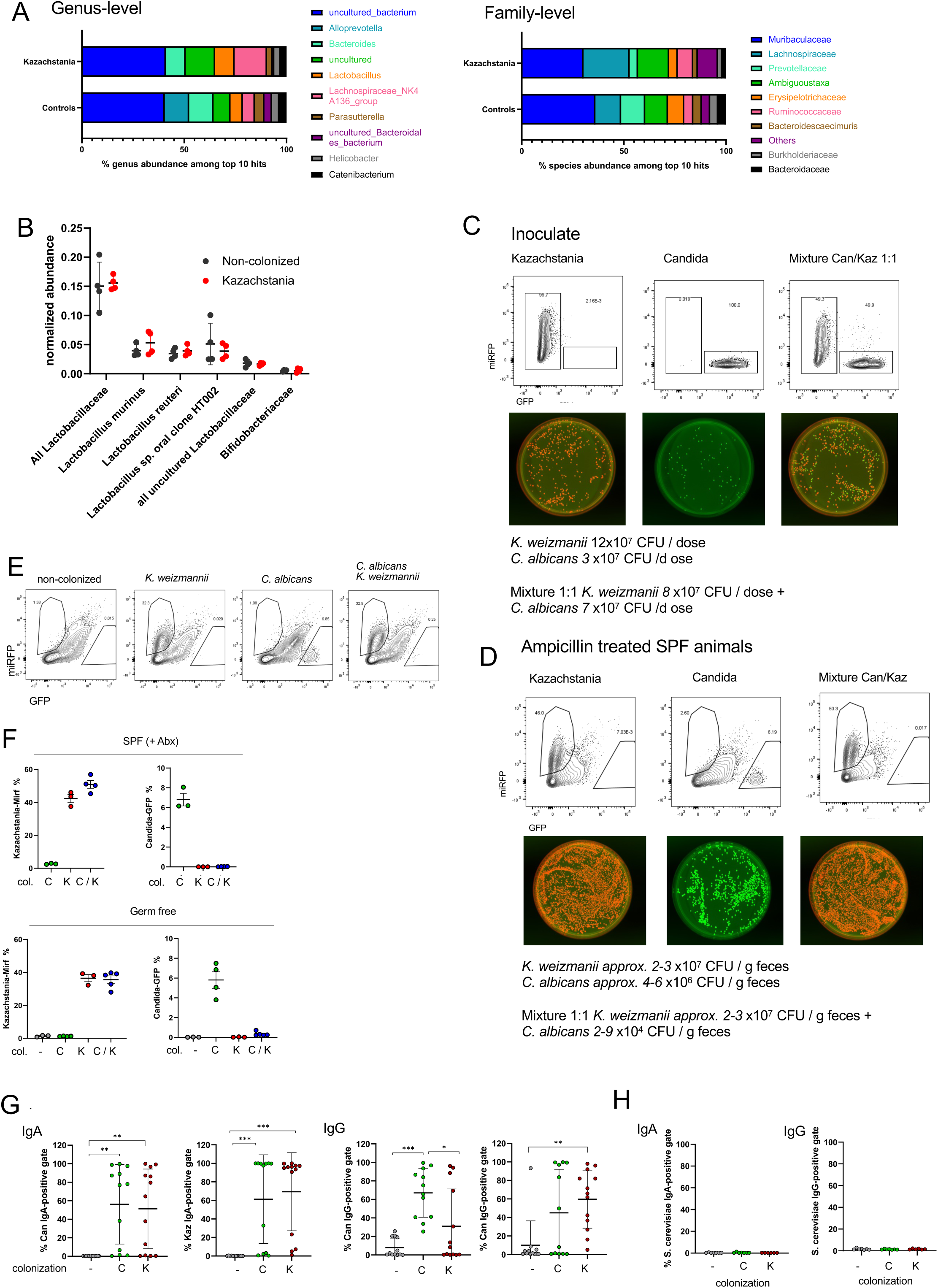
Impact of *K. weizmannii* colonization on microbiome. (A) Taxonomic composition of fecal bacterial identified in *K. weizmannii* colonized or non-colonized group (controls), as determined by 16S rRNA sequencing. The top 10 categories with the highest average relative abundance are shown at genus and family levels. (B) Normalized abundance of fecal *Lactobacillaceae* (divided into specific *Lactobacilli* species) and *Bifidobacteriaceae*, determined by 16S sequencing. (C, D) *In vivo* competition between *C. albicans* and *K. weizmannii* upon co-administration in Germ-free animals and in antibiotica treated wild type mice: FACS and cultivation analysis of culture inoculate for p.o. administration (C) Representative picture of recovery of C. albicans ENO1-GFP and *Kazachstania* ENO1-miRFP from feces of colonized animals using flow cytometry and cultivation (D) (E) Flow cytometric analysis of feces of germfree animals 1 week after oral inoculation of single fungi cultures or mixed *C. albicans* SC5314 (ENO1-GFP) / *K. weizmannii* (ENO1-miRFP) culture. (F) Percentages of *C. albicans* and *K. weizmannii* recovered from feces of germ-free animals colonized with *C. albicans* SC5314 (C), *K. weizmannii* (K) or mixed culture (C/K) compared to Abx-treated SPF animals 1 week after administration as determined by flow cytometry analysis (K); representative of 2 independent experiments. (G) Graph summarizing results serum reactivity of C*. albicans* or *K. weizmannii*-colonized animals (data shown in Fig. 4C). Y axis represents percentage of anti IgA or IgG-positive gate with mean ± SEM, N=11-13 mice per group. (H) Summary of flow cytometric analysis of humoral IgA and IgG anti-fungal reactivity for cultured *S. cerevisiae,* using sera for which anti-C*. albicans* or *K. weizmannii* reactivity was shown in Fig. 4C.

**Suppl. Figure 5:**
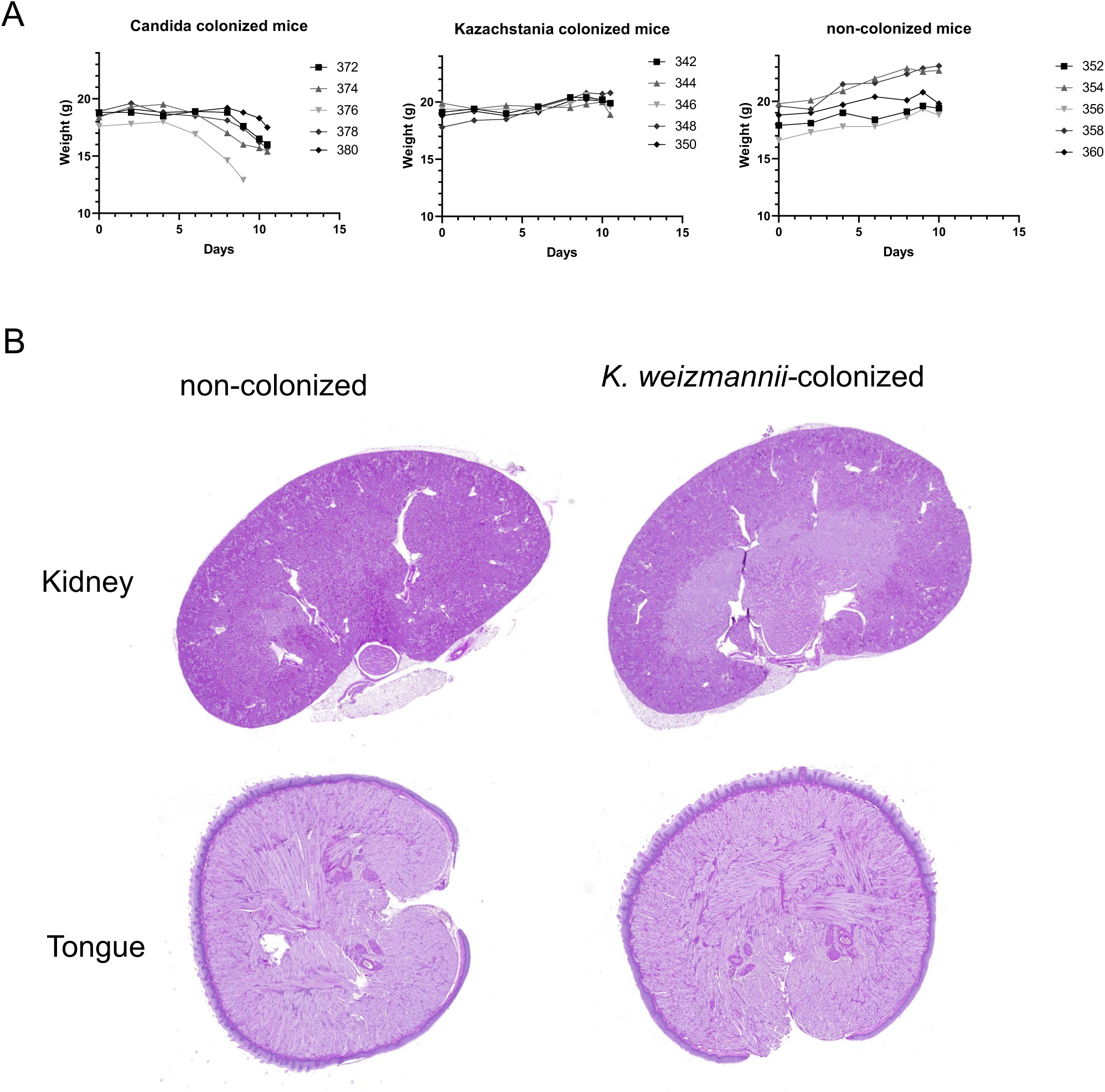
Analysis of *C. albicans-* colonized immunosuppressed animals. (A) Absolute weight monitoring (g) of individual mice in *C. albicans* colonization, *K. weizmannii* colonized and non-colonized group. group upon immunosuppressive treatment (B) Representative picture of kidneys and tongues (PAS staining) of immunosuppressed non-colonized and Kazachstania-colonized animals in end time point (day 10)

**Suppl. Figure 6:**
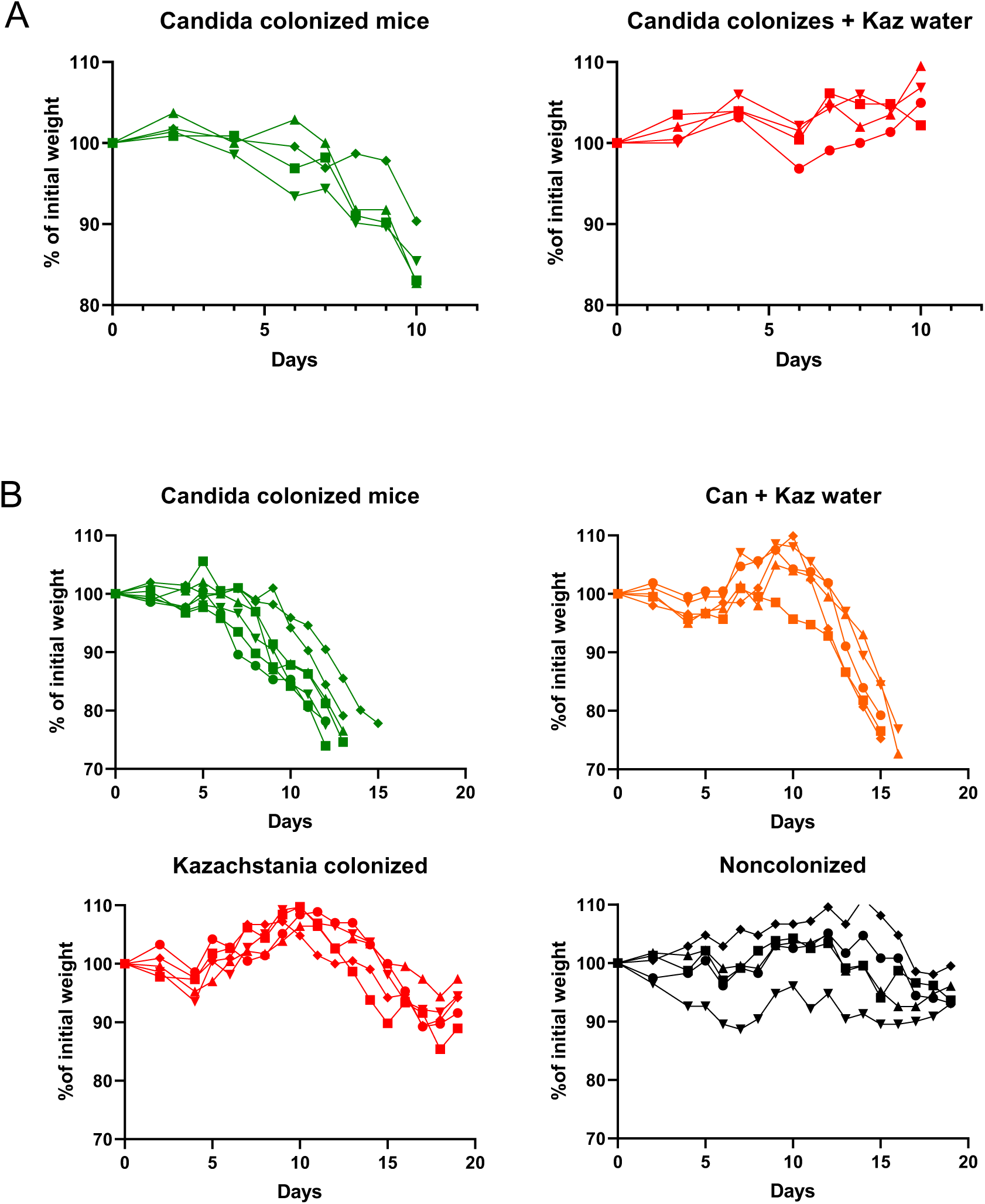
Analysis of *K. weizmannii* and *C. albicans-* colonized immunosuppressed animals. (A) Absolute weight (g) monitoring, comparison of immunosuppressed animals following *C. albicans*-colonization and *K. weizmannii*-outcompeted *C. albicans* colonization, monitoring till the end of experiment (day 10) (B) Absolute weight (g) monitoring showing the prolonged course of immunosuppressive treatment (mice sacrificed upon >20% weight loss), comparison of immunosuppressed animals following *C. albicans*-colonization and *K. weizmannii*-outcompeted *C. albicans* colonization, *K. weizmannii* colonized and non-colonized animals.

**Suppl. Figure 7:**
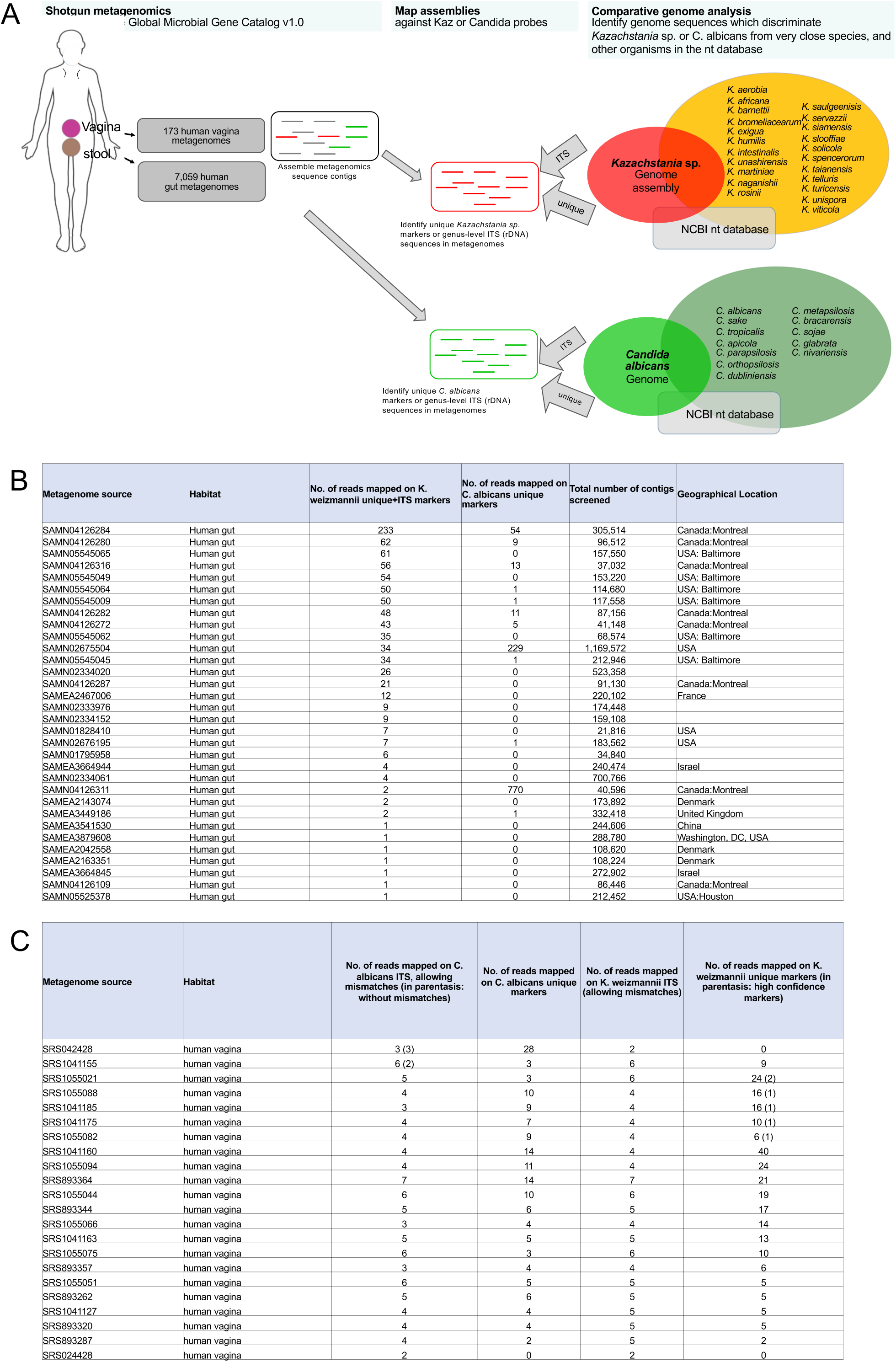
Human metagenomes identified with *K. weizmannii*. (A) Schematic of bioinformatic screening strategy of metagenomes. Briefly (from right to left), comparative genome analysis identified nucleotides regions that are unique to the genomes of either *K. weizmannii*. or *C. albicans* species, together with genus-level regions of their ITS sequences. These regions were used to screen thousands of shotgun metagenomics collected and assembled by the Global Microbial Gene Catalogue v1.0 (Coelho et al., 2022) (see methods) (B) Metadata and read counts of 32 metagenomes from human gut origin, in which *K. weizmannii* was identified. “Number of mapped reads” relates to assembled contigs of each metagenomic sample (provided by the Global Microbial Gene Catalog (Coelho et al., 2022)), which was used as a query and mapped against either *K. weizmannii* or *C. albicans* markers. The metadata information was collected from NCBI Sequence Read Archive (SRA) data. (C) Metadata and read counts of 22 metagenomes from human vaginal origin, in which either *K. weizmannii* or *C. albicans* were identified.

